# Quorum sensing underlies the establishment and maintenance of marginal zone B cells

**DOI:** 10.64898/2026.07.11.737964

**Authors:** Apoorva Singh, Melissa Verheijen, Thea Hogan, Andrew J. Yates, Benedict Seddon, Sanket Rane

## Abstract

The marginal zone (MZ) of the spleen harbors B cells that play an indispensable role in immune defense, orchestrating rapid responses to bloodborne pathogens. Under steady-state conditions, MZ B cell numbers are maintained through the balance of *de novo* generation from precursors, proliferative self-renewal, and loss. The mechanisms governing this homeostatic control remain elusive. Further, the developmental pathways underlying the establishment and continued supplementation of the MZ B cell compartment are not fullyelucidated. To address these gaps, we combined multiple fate-mapping tools and mathematical models to study MZ B cell dynamics in mice across the life course. Our analyses find evidence of quorum sensing mechanisms that regulate both the accumulation of mature MZ B cells during early life and their maintenance throughout adulthood. Specifically, we demonstrate that they derive predominantly from transitional B cell precursors with an efficiency that increases with age, reaching stable levels only in adulthood. MZ B cells compensate for this early developmental inefficiency through cell density-dependent proliferation, ensuring the timely establishment of a stable pool. Collectively, these findings unveil critical roles of quorum sensing and immune system maturation in the maintenance of this vital B cell subset.

## Introduction

Efficient resolution of systemic infection relies on the rapid production and secretion of antibodies. A critical component of this protection is mediated by marginal zone (MZ) B cells that continuously surveil the circulation through the spleen and respond to pathogens that emerge in blood ^6,19,31,46^. Beyond this frontline role, MZ B cells can initiate and augment the high affinity antibody responses mounted by follicular (Fo) B cells ^4,11,12,31,39^, and contribute to maintaining the natural antibody repertoire ^9,16,22,30^.

Consensus holds that fully mature Fo and MZ B cells derive from transitional B cells, transient populations that derive directly from immature B cells in the bone marrow ^2,29,34^. Studies using BrdU labeling and fate reporter systems have shown that in healthy adult mice Fo and MZ B cells cells persist on average for 5 weeks ^43^ and 3-7 months ^10,25^, respectively. To achieve stability in numbers, this loss through death or onward differentiation must therefore be balanced with compensatory influx and/or self-renewal. We have shown that Fo B cells self-renew rarely and are likely replenished largely through *de novo* generation from early-stage transitional (T1) B cells at rates that increase through life ^43^. The contributions of production and self renewal to the maintenance of MZ B cell numbers across the lifespan remain poorly defined.

An additional layer of uncertainty is the extent to which rates of immigration, division and/or loss are subject to regulation or feedback. In a study that blocked the production of B cells in the BM through conditional RAG2 deletion, MZ B cell numbers remained stable for over a year, whereas Fo B cell numbers gradually declined ^21^. This stability in the absence of influx is consistent with a long-standing view that MZ B cell numbers are determined not by the supply of precursors but by the carrying capacity of their anatomical niche. This idea originates from early work on B cell clonal selection into the MZ compartment, where the rate of clonal production and BCR-dependent survival signals were shown to determine enrichment in the CD21^high^ MZ compartment relative to the follicle ^29^, suggestive of clone-specific competition for resources. This idea was made explicit by later work proposing that the capacity of the splenic MZ niche to retain B cells is limited, such that B cells in excess of this capacity instead populate the follicle ^26^. A cellular and molecular basis for this niche was subsequently identified in the form of a population of DLL1-expressing stromal cells in the splenic red pulp, for which MZ precursors compete via Notch2 signalling. The strength of this competition — and hence the size of the resulting MZ B cell pool — is tuned by Fringe-dependent glycosylation of Notch2^42^. This model was further supported by competitive bone marrow chimera experiments showing that only B cells capable of efficient Notch2-DLL1 interaction are able to populate the MZ niche ^27^. It remains unclear, however, what cellular processes are modulated by niche occupancy, and how these may change with age; survival, onward differentiation, the ability to self-renew, or the ability to accommodate newly immigrated cells.

The immediate precursor of mature MZ B cells is also unclear. A prevailing view was that they derive from late-stage transitional (T2) B cells, via an intermediate marginal zone precursor (MZP) subset characterized by high expression of CD21 and CD23^41^. More recent work, however, has challenged this simple linear pathway by demonstrating that MZ B cells can differentiate directly from T1 B cells ^7,20,25^ or even from mature Fo B cells ^5,27^. Resolving the contributions of these pathways to MZ B cell homeostasis is critical for accurately modeling their population dynamics and predicting how they respond to physiological perturbations, including infections and aging.

To address all of these uncertainties, we employed an integrative approach combining mathematical modeling with a fate mapping mouse system ^23,35,43^. We characterized the kinetics governing the percolation of newly generated B cells into peripheral subsets and coupled it with measurements of proliferation (Ki67 expression), explicitly accounting for the role of self renewal in maintaining mature populations. We then extended our models to describe the kinetics underlying the establishment of MZ B cell pools during early life. Collectively, our analyses support a model in which competition for homeostatic resources governs MZ B cell population dynamics, and that they derive predominantly from transitional B cell subsets at a rate that increases with age.

## Results

### The marginal zone B cell compartment undergoes continuous replenishment by bone marrow-derived precursors

To elucidate the mechanisms regulating MZ B cell development and maintenance, we used a temporal fate mapping system to track the de novo production of MZ B cells across the life course ^24,43^. To do this, we generated chimeric mice by adoptively transferring T and B cell-depleted bone marrow (BM) cells from CD45.2 donor mice to reconstitute the hematopoietic system of CD45.1 recipient mice treated with the transplant conditioning agent busulfan (**Figure 1A**). Conditioning with optimized doses of busulfan selectively depletes the BM hematopoetic stem cell (HSC) population while leaving mature peripheral hematopoietic lineages intact. This procedure typically results in 60%–95% donor:host chimerism in the BM HSC compartment, which remains stable over the mouse lifetime ^17,23,24,43^. This system allows us to examine lymphocyte behavior under homeostatic conditions, without confounding side-effects typically observed in irradiation-based chimera models, and allows the visualization of the percolation of donor-derived cells into the mature peripheral B cell compartments without perturbing their numbers. Leveraging this approach, we followed the replacement of mature peripheral B cells by donor HSC-derived progeny for around 1 year post-BMT.

**Figure 1.**
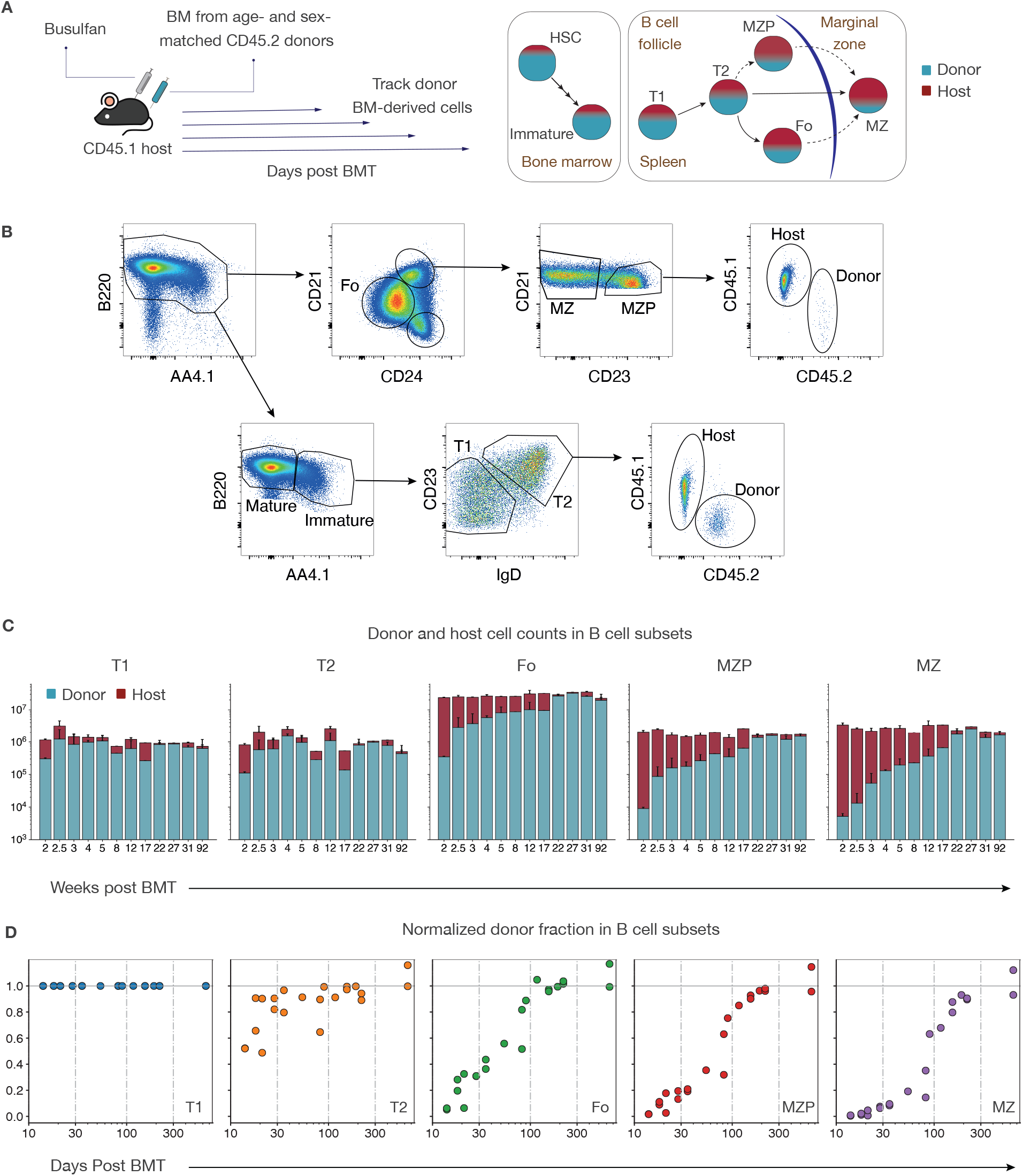
Tracking constitutive replenishment of B cell subsets using busulfan chimeras. **A**. Schematic of the busulfan chimera experiment. We also illustrate the likely path(s) taken by donor HSC-derived progeny as they progress through the B cell lineage, based on prior evidence, and how this may shape the composition of new donor bone marrow-derived cells versus pre-existing (host) cells across peripheral developing B cell stages. **B**. Gating strategy used to identify donor- and host-origin cells within splenic B cell subsets. Splenocytes from busulfan chimeras were stained with fluorescently labeled antibodies against B220, CD24, CD21, CD23, CD45.1, and CD45.2. **C**. Overall trends in donor-host composition and total pool sizes (mean + SEM) across the different B cell subsets in adult mice (n=22, ~2-24 months). Note that the x-axis represents a categorical variable, while the y-axis is plotted on a log10 scale. **D**. Donor fraction normalized to chimerism in T1 B cell subset across splenic B cell developmental stages

We studied donor cell influx into well-characterized developmental stages in the splenic B cell compartment using flow cytometry (**Figure 1B**). Developing B cells from different BM sites converge into the spleen as AA4.1^+^ IgM^high^ CD23^−^ transitional 1 (T1) cells. T1 B cells subsequently mature into IgD^high^ CD23^+^ transitional 2 (T2) B cells, which circulate between secondary lymphatic tissues. These transient subsets mature into IgD^high^ CD23^+^ CD21^int^ Fo and IgM^high^ CD23^−^ CD21^+^ MZ B cells ^2,29,34^. We partitioned total splenic B220^+^ B cells into three distinct subsets based on differential expression of CD24 and CD21 surface markers (**Figure 1B**). The CD21^high^ CD24^high^ subset encompasses both mature MZ B cells and CD23^high^ MZ B precursors (MZP) as described previously ^41^. We defined follicular (Fo) B cells by intermediate expression of both CD24 and CD21. Transitional B cells were pulled from the total B cell population based on their high expresion of the C-type transmembrane lectin CD93 (AA4.1). Within this population, T2 B cells were distinguished from T1 B cells based on their heightened CD23 and IgD expression. Lastly, we distinguished donor- and host-derived cells within each subset using CD45.1 and CD45.2 expression (**Figure 1B**).

The rates at which donor-derived cells accumulate in these subsets reflect their developmental sequence and their rates of replacement (**Figure 1C**). Consistent with their early developmental stage and transient nature, T1 and T2 subsets became rapidly enriched in donor cells. Donor cells progressively accumulated in more mature peripheral B cell subsets while the total size of each remained constant for nearly 2 years post-BMT. Therefore Fo, MZP and MZ cells all underwent continuous replacement from precursors during this period. Notably, at each time point and in every animal, the donor cell fraction was lowest within MZ B cells, indicating that on average they turn over more slowly than the other mature subsets. Collectively, these findings indicate that in adult mice, all of these subsets are continually replenished by bone marrow-derived cells.

### Donor infusion kinetics reveal of the rules replacement and precursor-progeny hierarchy across B cell developmental stages

Levels of HSC depletion and replacement varied across mice. Therefore, in order to describe the kinetics of donor chimerism (fraction donor-derived) in B cell subsets across animals we normalized the chimerism within each subset to that of a common developmental precursor in the same animal. We employed a similar strategy to study lymphocyte dynamics using busulfan chimeras in other settings ^17,23,43^. We have previously shown that chimerism in the splenic T1 compartment is a good proxy for the average HSC chimerism across all BM sites in a mouse ^43^, as all developing B cells enter the periphery via this transitional stage. For every downstream population, we defined its ‘normalized donor fraction’ to be

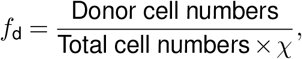

where χ is the the chimerism in T1.

The kinetics of *f*_d_ are rich in information. First, if *f*_d_ converges to 1 in a population, its chimerism mirrors that of its T1 B cell ancestors, and so it has turned over completely. By implication, then, donor cells (new) and host (which include cells generated pre-BMT) are equally replaceable. An asymptotic *f*_d_ *<* 1 indicates that replacement is incomplete. This can arise in several ways. The first is that influx has declined before the subset can turn over completely. It can also arise if host (older) cells are more persistent than donor (newer) cells, on average, either that subset or in its precursor. This fitness difference must derive from an increase in the rate of self-renewal and/or a decrease in the loss rate with time since a cells’ ancestor was generated in the BM. However, we saw that all naive B cell subsets exhibited *f*_d_ → 1 in the 90 weeks post-BMT (**Figure 1D**), indicating that replacement in all of these subsets was continuous and that there was no detectable signal of dynamics varying with cell age.

Second, the kinetics of *f*_d_ in different subsets also contain information regarding their relative positioning within the developmental hierarchy. Assuming host and donor cells are equally able to differentiate, the rate of approach to a stable *f*_d_ within a subset cannot be faster than that of its immediate precursor. Since *f*_d_ increased most slowly in MZ B cells, some or all of the other subsets T1, T2, Fo, and MZP may serve as their potential precursors.

Third, the rate at which *f*_d_ stabilizes in a subset reflects the rate of influx from a precursor population. If the subset remains at a constant size, this rate of influx must equal the net rate of loss – the balance of selfrenewal and loss through death, and onward differentiation. Thus, using mathematical models of the kinetics of donor cell replacement across these subsets allows us to build a detailed, quantitative picture of their relationships and dynamics.

### Ki67 expression as a readout of self-renewal and ancestral division in mature B cell subsets

To examine the role of cell division in sustaining MZ B cell numbers we measured levels of Ki67, a nuclear protein that is expressed upon entry into the G1 stage of cell cycle and remains detectable for several days postmitosis ^32^. Ki67 expression within a population therefore provides a readout of its recent division history. Paired with the kinetics of donor cell replacement, Ki67 allows us to separate the net rate of loss into self-renewal and death/differentiation. We have also shown that high levels of Ki67 expression on T1 and T2 cells of the spleen are likely predominantly residual protein from the division activity of their precursors in the bone marrow ^43^, rather than a sign of self-renewal.

We saw clear unimodal patterns of Ki67 expression within Fo, MZP, and MZ B cells, with 15-22% of cells being Ki67^+^ and MZ B cells at the upper end of this range (**Figure 2A**). Taken at face value, these proportions indicate substantial levels of division within these subsets. However, given the short transit times through the T1 and T2 stages, which express Ki67 at high levels, it is plausible that at least some of this expression is a continuation of this ‘shadow’ of earlier divisions within highly proliferative BM precursors, rather than self-renewal alone. Consistent with this idea, we observed that the distributions of Ki67 expression among Fo, MZP and MZ B cells overlapped with the tail of the distribution within T2 B cells, whose distribution in turn overlapped that of T1 B cells (**Figure 2A**). These patterns of expression remained roughly constant with time (**Figure 2B**), with a suggestion of a decline in the Ki67^+^ fraction within Fo B cells.

**Figure 2.**
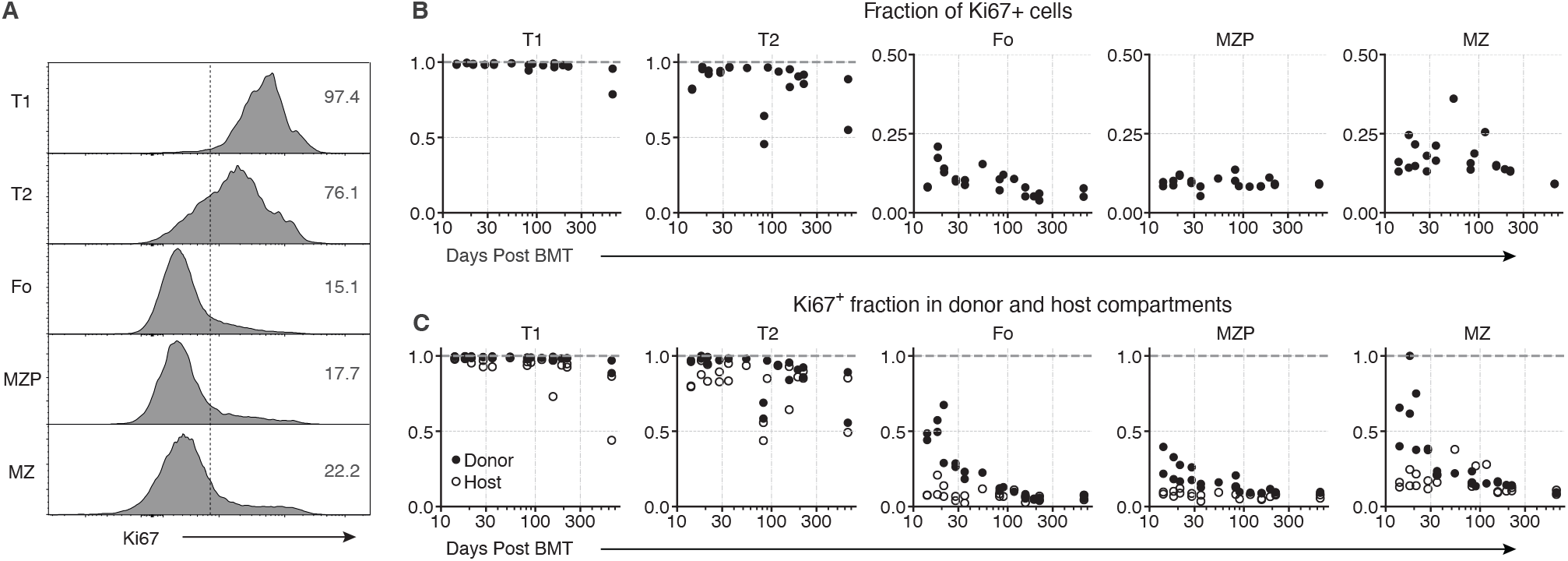
Ki67 expression dynamics throughout B cell development. **A**. We show a representative plot of Ki67 expression profiles across B cell developmental stages in the spleen. Numbers in each panel indicate % of Ki67^+^ cells within each subset. **B**. The proportions of Ki67-expressing cells within the peripheral B cell developmental stages. **C**. The proportions of Ki67-expressing cells within the donor and host sub-compartments for splenic B cell subsets.

Next, we analyzed the proportion of Ki67^+^ cells within the donor and host compartments separately, across B cell subsets and over time (**Figure 2C**). The Ki67^+^ fraction remained consistently elevated and comparable in both donor and host transitional subsets, consistent with their transient status (**Figure 2C**). Mature B cells presented a contrasting pattern of expression. Shortly after BMT, the donor Fo, MZP, and MZ B cells had higher Ki67 fractions compared to their respective host cells. These elevated fractions subsequently decreased to host levels after roughly 3 months. The simplest explanation of this dynamic does not invoke any intrinsic differences between host and donor cell behavior; rather, it derives from short-lived differences in their recent histories. In the first days to weeks after BMT, donor cells within a subset must predominantly be recent immigrants. Their Ki67 levels will be transiently elevated relative to established host cells if they derive directly from a Ki67-rich precursor, such as T1 or T2, or if their differentiation into that subset is associated with cell division. Over time, the residence profiles of host and donor cells (i.e. their distributions of times since entry into the compartment) converge, along with their mean levels of Ki67 expression. This transient behavior following BMT is highly informative regarding B cell subset dynamics. Therefore, in models described below, we allow for Ki67^+^ expression to derive both from inheritance and self-renewal.

### Describing MZ B cell dynamics in adult mice with a simple model of homogeneous turnover

In order to use these data to probe the ontogeny and homeostatic regulation of MZ B cells, we developed a mechanistic modeling framework in which the population dynamics of putative precursor populations serve as explicit constraints on predicted MZ B cell behavior (**Figure 3A**). Our goal was to infer the model parameters to optimally recapitulate the observed kinetics of *f*_d_ and Ki67^+^ frequencies in MZ B cells for each candidate precursor population. We evaluated T1, T2, MZP, and Fo B cells independently as potential dominant precursors to the MZ B cell compartment.

**Figure 3.**
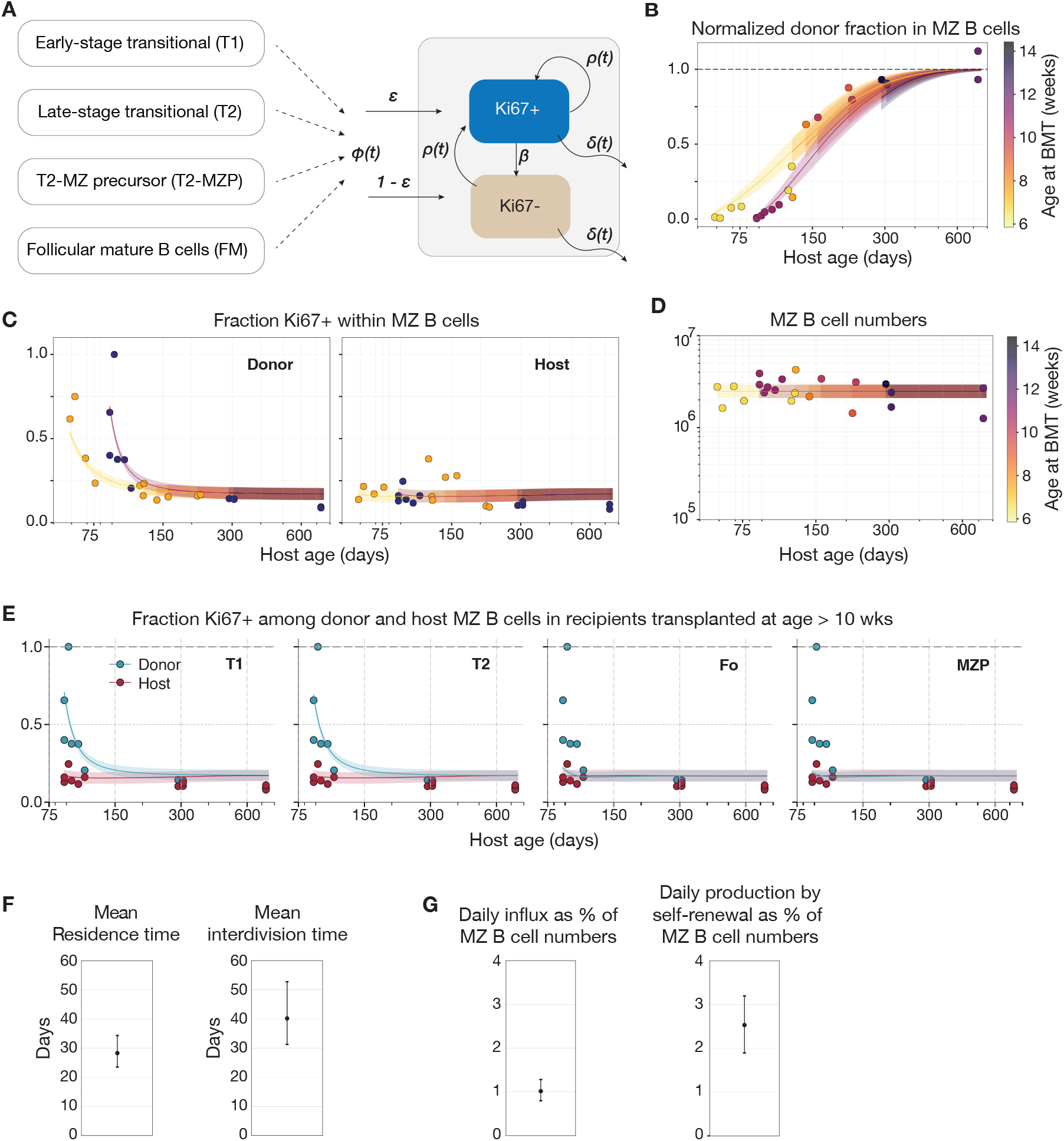
Modeling the population dynamics of MZ B cells in adult busulfan chimeric mice. **A**. We depict a general formulation encompassing all candidate models of MZ B cell dynamics, in which the rates are either held constant or allowed to vary with time. In the constant birth–loss model, *ϕ*(*t*) = *ϕ, ρ*(*t*) = *ρ*, and *δ*(*t*) = *δ*. In the time-varying model, *ϕ*(*t*), *ρ*(*t*), and *δ*(*t*) are instead treated as time-varying functions (see **SI section S2** for details). **B–D**. Predictions from the constant birth-loss model considering both T1 (dashed) and T2 (solid) as precursors, shown for (B) normalized chimerism (relative to the T1 compartment), (C) the fractions of donor (first subplot) and host (second subplot) cells that expressed Ki67, and (D) MZ B cell counts. **E**. Ki67^+^ frequencies among donor and host MZ B cells in older recipients (BMT after 10 weeks of age) across all candidate precursors, illustrating the discrepancy between model predictions and observed Ki67 kinetics when Fo B or MZP are assumed as precursors. **F and G**. Parameters inferred from the analysis of busulfan chimera data using the constant birth-loss model (22 mice *≥*7 weeks). (F.) Estimated mean residence and inter-division times of MZ B cells with 95% credible intervals. (G.) The left panel depicts the daily influx of T2 B cells into MZ B compartment expressed as a percentage of the MZ B cell pool size. In the right panel, we show the contribution of self renewing divisions to the MZ B cell maintenance, expressed as a percentage of the MZ B cell pool size.

We began with the simplest model of constant birth and loss in which new MZ B cells — whether of host or donor origin — enter the compartment at a total rate proportional to the size of their precursor population (*P* (*t*)), *ϕP* (*t*). Existing MZ B cells self-renew through division at *per capita* rate *ρ* and leave the population through death or onward differentiation at *per capita* rate *δ* (**Equation 1** and **Figure 3A**). Since MZ B cell numbers (*M*) remain stable in adult mice (**Figure 1C**), we imposed a steady-state assumption in which production is balanced by loss:

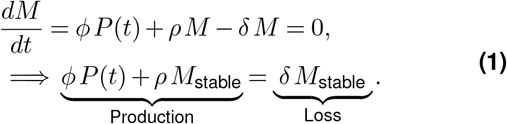

The precursor population was taken to be one of T1, T2, Fo, or MZP B cells. We duplicated this model for host and donor cells, with differences deriving only from the time-dependent chimerism within the precursor, which, together with its total population size *P*, we described empirically (**SI section S1** and **Figure S1**). Finally, we stratified host and donor cells each into Ki67^+^ and Ki67^--^ cells, allowing for inheritance from the precursor by explicitly modeling the Ki67 content of immigrating cells, again with empirical functions (**SI section S1**).

Our data derived from 22 mice that were aged between 7-15 weeks at transplantation. Therefore data from each animal derived from a potentially different ‘history’ of *de novo* generation from the precursor. We accounted for any effect of this variation by incorporating the recipient age at BMT as a variable within the modeling framework. Doing this allowed us to generate prediction curves for animals of varying ages using a shared parameter set. The formulation of the models describing the dynamics of Ki67^+^ and Ki67^--^ subpopulations within the donor and host MZ B cells is described in **SI section S2**.

We employed a Bayesian statistical framework to estimate model parameters and to quantify uncertainty in model predictions. Candidate models were compared using leave-one-out (LOO) cross-validation, which yields the expected log pointwise predictive density (*ELPD*) as a measure of predictive performance (**SI section S3**). Each precursor:model combination was fitted simultaneously to four time courses — MZ B cell numbers; the donor chimerism normalized to that of T1 cells (the earliest common precursor among all populations considered), *i*.*e. f*_d_; and the proportions of Ki67^+^ cells among host and donor MZ B cells. Full details of the fitting procedure and model selection strategy are provided in **SI section S3**.

### MZ B cells are continually replenished from transitional populations and undergo near complete replacement yearly

Using this approach we found the strongest and approximately equivalent support for transitional T1 and/or T2 B cells as the dominant precursors of MZ B cells. For both T1 and T2 as precursors, the simple constant birth-loss model produced smooth *f*_d_ trajectories approaching a stable plateau consistent with eventual complete donor replacement across recipient age groups (denoted by different colors in **Figure 3B**). It also captured the trajectories of the Ki67^+^ fractions among host and donor MZ B cells (**Figure 3C**) and steady-state population size across animals (**Figure 3D**). Models with Fo or MZP B cells received very little support (**Table 1**), largely because neither expressed Ki67 themselves at sufficiently high levels to explain its strong enrichment in donor cells immediately following BMT (**Figure 3E**).

**Table 1.**
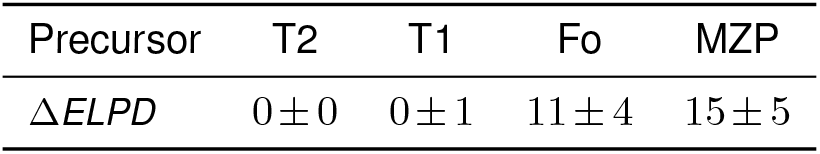
Support for each candidate precursor under the constant birth-loss model. We use the *ELPD* estimate as a measure of the goodness-of-fit, where the precursor with the highest *ELPD* is considered the best fit. For each precursor, we report Δ*ELPD*, the absolute difference between its *ELPD* estimate and the highest *ELPD*, together with the standard error (*s*.*e*.) of this difference. By convention, models whose Δ*ELPD* values fall within twice the *s*.*e*. are considered to have equivocal support.

While these fits looked reasonable, we wanted to assess whether host age influences MZ B cell dynamics, motivated by our previous finding that Fo B cells are lost more slowly as mice age ^43^. We therefore extended the model in two ways; one in which the percapita rate of recruitment from the precursor (*ϕ*) varies with age, and another in which the rate of self renewal (*ρ*) varies (illustrated in **Figure 3A**). In both models, the loss rate was allowed to covary with age to preserve the steady-state constraint, as described in **SI section S2**. Neither extension improved model fit nor received greater support from the data relative to the constant birth-loss model (**SI able S1**). Moreover, allowing for age-dependent dynamics did not alter the ranking of precursor populations. Across all model variants, T1 and T2 subsets consistently received the strongest support. The **SI figure S2** displays all alternative model fit results, while **SI section S2** contains parameter definitions for all models.

Parameters estimated using either T1 or T2 as precursors were comparable (**SI table S2**). In **Figure 3F**, we present parameters estimated using T2 as precursors, for consistency with the prevailing view in the field. We estimated the mean residence time of MZ B cells – the average time for an MZ B cell to leave the compartment by death, egress, or differentiation – to be approximately 28 days, with inter-division times of around 40 days. This indicates that MZ B cells are slightly more likely to exit the compartment, through death or onward differentiation, than to divide within it. Based on these estimates, we infer that the MZ B cell population is replaced almost completely within approximately 1 year (see **SI section S4**). We also estimated that influx from transitional precursors replaces roughly 1% of the MZ B cell compartment daily, while self-renewal contributes ~2.5% of the pool each day (**Figure 3G**). These inferences support the notion that Ki67 levels observed in MZ B cells reflect a combination of inheritance from precursors and proliferation within the compartment itself. This profile contrasts sharply with Fo B cells, who self-renew rarely and whose Ki67 is almost entirely inherited from precursors ^43^, highlighting distinct homeostatic mechanisms in these two mature B cell subsets.

### Early life dynamics of MZ B cell development differ from those in adults and involve quorum-sensing

It is unclear whether the ontogeny and dynamics of MZ B cells that we inferred in adults mice apply in the neonatal period, when peripheral lymphocyte subsets have yet to reach steady state. To explore this, we analyzed T1, T2, and MZ B cell numbers in young mice over the first 20 weeks of life. We took advantage of data from experiments analyzing Rag2-EGFP Ki67-RFP reporter mice, described previously ^43^. After birth, numbers of both T1 and T2 B cells rose sharply up to approximately 3 weeks, then declined over the following ~3 weeks and were thereafter maintained at a stable level (**Figure 4A**). MZ B cell numbers increased up to ~8 weeks before stabilizing at the levels we observed in adults (**Figure 4B**).

**Figure 4.**
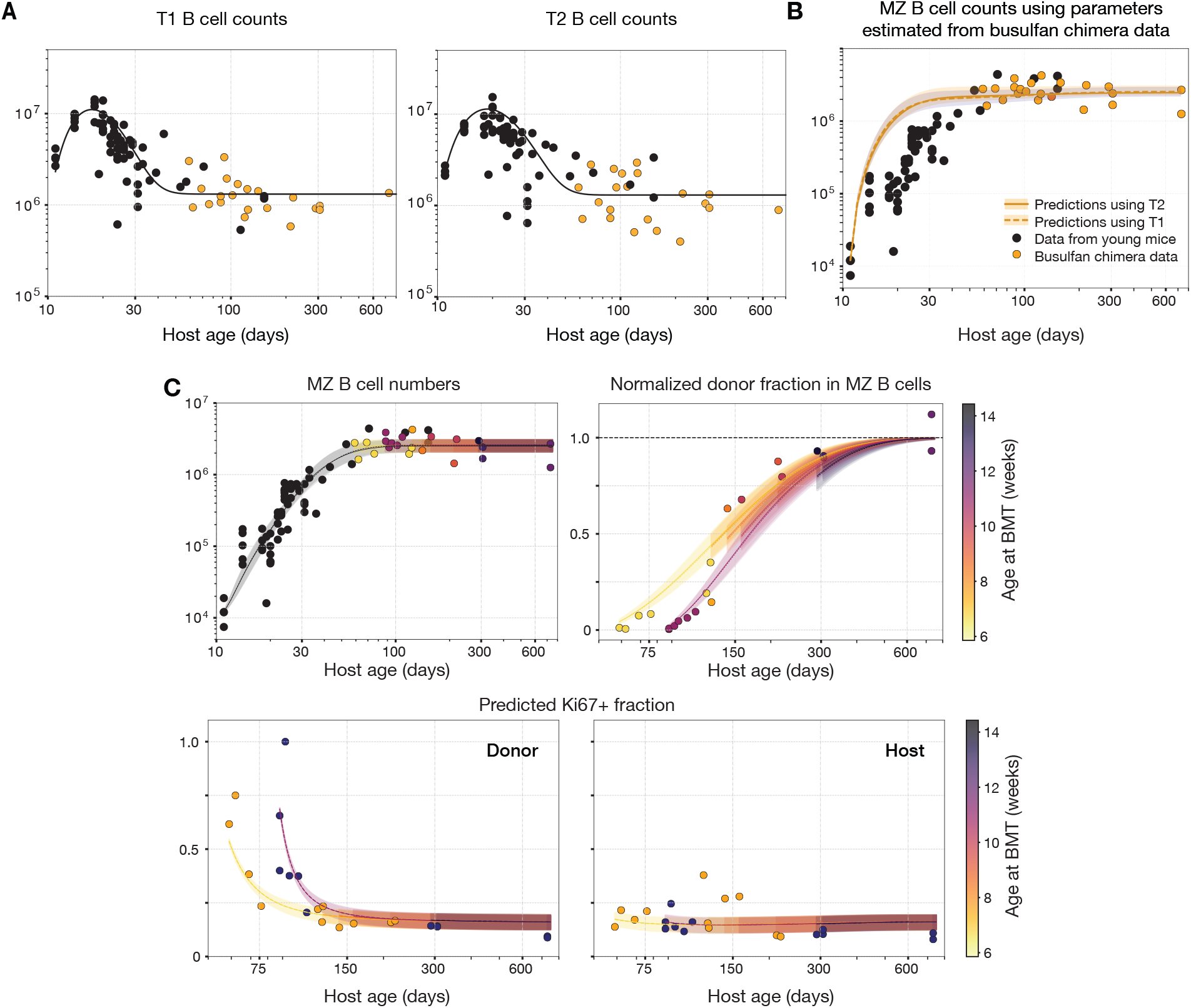
MZ B cell establishment dynamics in early life. **A**. Numbers of splenic T1 and T2 B cells recovered from both young WT mice (n = 62, black) and adult busulfan chimeras (n = 22, orange), described with a single fitted empirical function (see SI section S5) with a black line showing the mean. **B**. MZ B cell numbers in young WT mice (n = 62, black) and busulfan chimeras (n = 22, orange). The black line shows the mean with 95% credible intervals from a prediction of the constant birth-loss model with T1 (dashed) and T2 (solid) as precursors, extrapolated to early life using the fitted empirical function for their respective dynamics. **C**. Fits of the density-dependent division model with time-varying influx to the MZ B cell numbers in both young WT mice and adult chimeras, their donor fractions normalized to chimerism in T1 B cells, and Ki67^+^ frequencies among donor and host cells in chimeric mice.

We first asked whether the constant birth-loss model, with either T1 or T2 B cells as precursors, could account for the establishment of the MZ B cell compartment during early life. To explore this we described the dynamics of transitional subsets in young mice with fitted empirical functions (see **SI section S5; Figure 4A**). We then used these with the parameter estimates derived from adult busulfan chimera data to predict MZ B cell accumulation in young mice from day 11 onward. These projections failed to recapitulate the observed kinetics of MZ B cell development, overestimating the rate of early accumulation (**Figure 4B**). This discrepancy clearly suggested that the rules governing MZ B cell development in early life are distinct from those operating in adulthood.

We then extended the model to allow processes controlling MZ B cell dynamics to vary as mice age; specifically, (i) the rate of precursor differentiation, *ϕ*; (ii) the MZ B cell loss rate, *δ*; or (iii) the rate of self renewal, *ρ*. We fitted each model simultaneously to the earlylife and busulfan chimera data, using transitional subsets as precursors and a single function describing their trajectories from soon after birth into late adulthood. This ‘global’ approach allowed us to leverage the constraints afforded by the adult chimera data to help constrain model behavior in young mice (see **SI section S5** for modeling details). However, all of these models achieved only modest improvements in predicting neonatal MZ B dynamics, as they still failed to capture the initial growth rate and the approach to steady state (**SI figure S3**).

Historically, the concept of a niche is ubiquitous in immunology. Feedback regulation is not an absolute requirement for the establishment of a steady state, but it is required to explain the preservation of a steady state under persistent perturbations, such as the apparent maintenance of MZ numbers when replenishment from precursors is restricted^21^. Given this, and the limitations of the quite flexible models of mouseage-dependent kinetics, we reasoned that quorum sensing mechanisms might underlie the distinct dynamics observed in neonatal and adult mice. Therefore, we also tested a variety of models that allowed rates of influx, self renewal, or loss to be modulated by the MZ B cell numbers. However, none of these modifications in isolation could explain the trajectories of MZ B cell numbers early in life (**SI figure S4**).

We inferred that both production and self-renewal processes make substantial contributions to MZ B cell maintenance in adults. Therefore, we asked whether the dynamics in neonatal mice could result from time or density-dependent effects on multiple processes at once. To explore this possibility in an unbiased manner, we systematically tested models in which all process rates could vary with either mouse age or population size, both independently or in combination. Strikingly, a model in which division is governed by density-dependent regulation, influx is time-varying, and the loss rate remains constant, was uniquely able to account for early life dynamics of MZ B cells. This model produced the closest descriptions of MZ B cell development and maintenance across the entire mouse-lifespan (**Figure 4C**) and received strongest support from the data (**SI table S3**). Parameter estimates from the best-fit model are provided in **SI table S4**.

### Age and cell density shape MZ B cell production early in life

The best fitting model indicated that cell production, rather than survival, was modulated most strongly during the establishment of the MZ B cell pool. The influx parameter *ϕ* reflects the efficiency with which precursors mature into MZ B cells, and we estimated that it increases roughly 10-fold from birth until stabilization at roughly 8 weeks of age (**Figure 5A**). This trajectory results in a low rate of *de novo* production of MZ B cells in neonates, despite the abundant supply of transitional B cells (**Figure 4A**). The rate at which MZ B cells self-renew follows the opposite trend (**Figure 5B**); we estimate they divide every 10 days (95% CI: 7–16) at age 2 weeks, when the population is sparse, slowing to every 42 (34–50) days at age 10 weeks, when the compartment approaches carrying capacity (**Figure 5C**). In contrast we found no evidence for age or density dependence effects on MZ B cell lifespan (residence time), which remained at 30 days (25–34; **Figure 5C**). Strikingly, at steady state, the net loss rate of MZ B cells (the difference between loss and division rates) falls low enough that we estimate individual clones to have an average lifespan of about 100 days (CI: 85–125).

**Figure 5.**
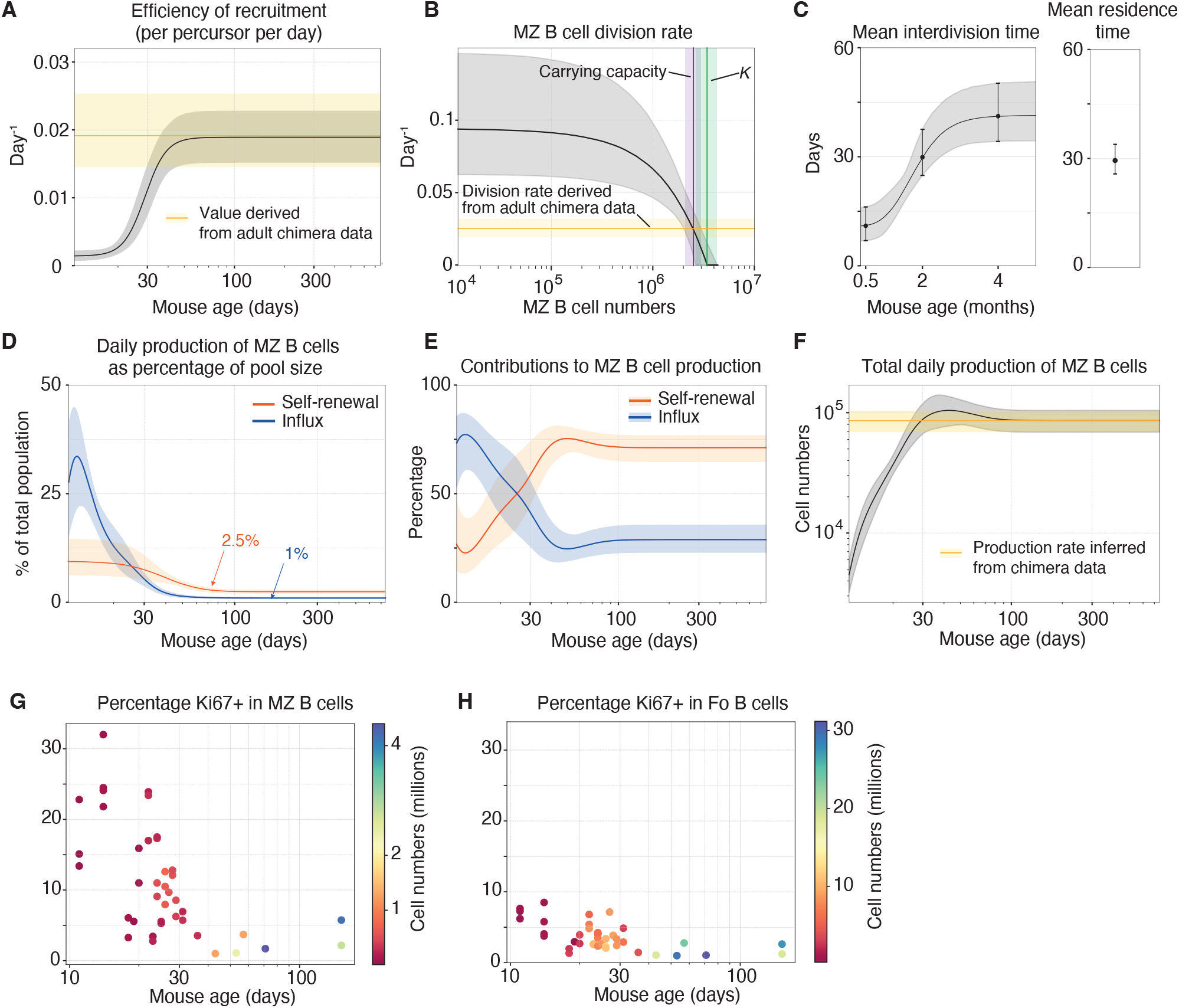
Insights into MZ B cell dynamics across development and homeostasis. Parameters estimated by simultaneously fitting the combined model, which accounts for both the density-dependent division with time-varying influx, to early life and chimera datasets, using the T2 subset as the precursor population. Parameter estimates obtained using the T1 as the precursor are shown in the **SI Table S4** and are qualitatively similar. In all plots, mean estimates are represented by lines and solid dots; envelopes and error bars depict estimation uncertainty as 95% credible intervals. **(A)** The time-varying efficiency of recruitment of new MZ B cells (*ϕ*(*t*)) during development, which stabilizes to the value estimated from adult data. **(B)** The estimated division rate as a function of population size. This curve intersects the rate inferred from adult data at the stable MZ B cell pool size (*i*.*e*. the carrying capacity) and approaches zero as population size nears the crowding parameter *K*, which dictates the degree of density-dependence (see **SI section S7** for details). **(C)** Left panel: Mean interdivision time of MZ B cells (the inverse of the division rate, *ρ*(*t*)) as a function of mouse age. Right panel: The expected residence time (the inverse of the loss rate) of MZ B cells. **(D)** Daily contributions of self-renewal and influx from precursors as percentages of MZ B cell numbers, plotted as a function of mouse age. **(E)** Relative contributions of self-renewal and influx to MZ B cell production, with mouse age. **(F)** Total daily production (influx plus self-renewal) of MZ B cells across the mouse lifespan. **(G-H)** Observed fractions of Ki67^+^ cells within the MZ (G) and Fo (H) B compartments in RAG-eGFP Ki67-RFP mice, plotted as a function of mouse age. Colors denote total population sizes.

When represented as a fraction of the MZ B pool, total production through self-renewal also declines with age, replenishing roughly 10% of cells daily at 2 weeks but only about 2.5% in adulthood (**Figure 5D**), in line with our estimates derived from adult data alone. The daily influx from precursors expressed as a fraction of the MZ B pool follows a similar trajectory, peaking sharply around 2 weeks (~30% per day) before dropping off rapidly and settling near 1% per day by 8 weeks (**Figure 5D**). Collectively, these dynamics create a clear developmental shift in the modality **(Figure 5E**) and total rate **(Figure 5F**) of production of MZ B cells (see **SI section S6**).

Together, the combined push-pull effects of mouse age on influx and cell density on self-renewal are required to correct the overshoot in early MZ B cell production predicted by the constant birth-loss model (**Figure 4B**) and describe the approach to stability. For instance, rising influx alone cannot account for both the rapid compartment growth seen in the first few weeks and its subsequent sharp transition to a stable plateau. Density-dependent division supplies the missing piece, providing the most parsimonious mechanistic explanation. Our model extends the classic logistic growth framework ^44^, with carrying capacity emerging from the interplay of time-varying immigration, density-dependent division, and ongoing loss (see **SI section S7** for details).

To test the prediction of a higher rate of self-renewal early in life, we analyzed Ki67 expression in MZ B cells from young Rag-eGFP Ki67-RFP mice and indeed found that Ki67^+^ frequencies were the highest in the youngest mice and declined with age (**Figure 5G**). Further, Ki67^+^ fractions were markedly lower among Fo B cells than MZ B cells, even though both populations receive input from transitional precursors (**Figure 5H**), suggesting that – as in adult mice – self-renewal plays much smaller role in Fo B cell maintenance.

In summary, we show that MZ B cell development and maintenance are governed by the interplay of two regulatory mechanisms. First, we inferred that there is an approximately 10-fold increase in the rate of differentiation of transitional B cell precursors into MZ B cells over the first 8 weeks of life. We previously observed a similar pattern in Fo B cell development ^43^, suggesting a broader developmental principle. Second, we inferred density dependence in the rate of self-renewal of MZ B cells, such that they divide more rapidly at low population densities. Such quorum-sensing behavior is suggestive of competitive regulation of signals required for proliferation and may act to compensate for the relatively inefficient recruitment into the MZ B cell compartment early in life.

## Discussion

By combining experimental data with mechanistic mathematical modeling, we characterized the dynamics and maintenance of marginal zone (MZ) B cells throughout the mouse lifespan. Our findings reveal that MZ B cells are continuously replenished by bone marrow-derived precursors, with a constant rate of replacement throughout adulthood.

The dynamics governing MZ B cell establishment in early life, however, differ markedly from those in adulthood. As well as age-dependent increases in the efficiency of differentiation into MZ B cells, we saw strong evidence of cell density-dependent regulation. Specifically, the *per capita* division rate of MZ B cells declines as their numbers increase, indicating elevated proliferation when the MZ niche is sparsely populated and a progressive slowdown as the compartment approaches steady state. This behavior is consistent with a quorum sensing mechanism, in which competition for limiting niche resources increases as the compartment grows ^15,33^.

Our inferences regarding MZ B cell ontogeny align with previous studies identifying transitional B cells as their direct precursor ^1,2,8,28,41^, and with more recent evidence suggesting that commitment to the MZ fate can occur as early as the T1 stage ^7,20,38^. However, the poor performance of models with Fo or MZP cells as the dominant precursors does not exclude the possibility that these populations harbor committed intermediates in the differentiation pathway. We reconcile this with historical support for the MZP by arguing that the subset likely represents a heterogeneous mixture of immature and mature cells, and thus its dynamics may not agree with that required by the bona fide MZ B cell precursor. Likewise, although Fo-to-MZ transdifferentiation is documented under acute stimulation (e.g., immune activation or constitutive Notch2 signaling) ^5,27^, we found no evidence that this pathway plays a significant role in baseline MZ B cell homeostasis.

The widely accepted view that the MZ B cell compartment is a long-lived, self-renewing population stems primarily from observations showing that its size remains stable for over a year following the cessation of BM-derived influx via inducible RAG2 deletion ^21^. A quorum sensing model provides an alternative explanation for this stability; that when precursor influx is absent, the system compensates either through increased proliferation or reduced cellular loss. Our analyses suggest that density-dependent division is the dominant mechanism, at least in early life, but a density-dependent increase in cell survival remains a possibility. Under identical conditions, the Fo B cell compartment halves in size every 4 months, indicating that this regulatory mechanism specifically operates on MZ B cells. Investigating division and loss dynamics in the MZ B cell compartment under conditions ranging from partial depletion to oversaturation in future studies will help us disentangle the contributions of these two processes and yield a clearer picture of the mechanisms underlying MZ B cell homeostasis.

Prior estimates of MZ B cell lifespan have varied considerably across studies. Jones *et al*. estimated a mean lifespan of ~7 months from the decay kinetics of labeled MZ B cells (half-life ≈5 months) ^25^, while Chappaz *et al*. reported ~3 months using BrdU pulse-chase labeling ^10^. A confounding factor in such labeling studies is continuous precursor influx, which can replenish the labeled pool over time and thereby inflate lifespan estimates ^3,14,18^. Accurate inferences regarding cell lifespans require considerations of population structure — whether the compartment is homogeneous or composed of sub-populations with distinct dynamics — as well as potential influences of cell age and cell density on loss rates ^3,13,15,36,37^. Under the assumption of homogeneity we estimated the expected lifespan of an MZ B cell (or the mean time until its death, egress from the marginal zone, or differentiation) to be approximately 30 days. However, the process of self-renewal allows for the persistence of MZ B cells at a clonal level; using our estimate of their division rate at steady state, we estimate that MZ B cell clones halve in size every ~70 days. This figure is consistent with data reported by Chappaz *et al*. ^10^ in which the frequency of labeled cohort of MZ B cells fell by 50% in approximately 60 days.

Our models do not unequivocally identify either T1 or T2 as the dominant MZ B cell precursor population. Nevertheless, they reveal a developmental program in which the *per capita* rate at which precursors differentiate into MZ B cells (our parameter *ϕ*, or the ‘efficiency of recruitment’; **Figure 5A**) increases as the immune system matures. We estimate that the total rate of production of new MZ B cells from precursors rises from approximately 3500 cells per day in neonates to a stable 25000 cells per day in adults. We previously identified a similar age-dependent pattern for follicular B cells, whose efficiency of generation from T1 precursors increases during early development ^43^. Collectively, these findings suggest a global pattern of progressive B cell niche maturation in early life.

In summary, by combining multiple fate-mapping systems and mathematical models it is possible to dissect and measure the roles of precursor influx and self-renewal in the maintenance of MZ B cells, and resolve how these processes are shaped by age and population density. In doing so, our results reveal a dynamic regulatory regime in which population stability emerges from continuous competition for limiting resources.

## Methods

### Busulfan chimeras

Bone marrow chimeric mice were generated according to established protocols ^24^. C57Bl6/J mice served as bone marrow donors, and SJL.C57Bl6/J congenic mice were used as hosts. Host age ranges are specified in the figures. Age-matched donor bone marrow was obtained from femurs of C57Bl6/J mice. The donor marrow suspension was then subjected to immuno-magnetic selection to deplete T cells and B cells using biotinylated antibodies targeting CD3 (eBioscience, 1/500 dilution), TCR-*β*(eBioscience, 1/500 dilution), and B220 (eBioscience, 1/200 dilution). The captured immune complexes were bound to streptavidin-coupled Dynabeads (Life Technologies); the unbound fraction was subsequently purified, achieving depletion of mature T cells and B cells. Twenty-four hours following the final busulfan injection, a total of 8–10 million cells were injected intravenously (i.v.) into the recipient host mice. At designated time points post-BMT, host animals were euthanized and their spleens were harvested for subsequent analysis.

### Reporter mice

The mKi67mcherry-CreERT2 mice were generated through targeted replacement of terminal exon 14 within the Mki67 locus. This genomic modification involved inserting an upstream FRT-flanked neomycin cassette, followed by a downstream sequence comprising the mCherry fusion construct, IRES sequence, and CreERT2 cDNA. The initial mouse line was subsequently crossed with actin-FLPe mice to excise the neomycin selection cassette, followed by crossing into the Rosa26RYFP strain ^40^ to yield the stable mKi67mcherry-CreERT2 Rosa26RYFP double reporter line. Cre recombinase activity was induced in vivo in these animals via intraperitoneal (i.p.) injection of 2 mg of tamoxifen (Sigma) diluted in corn oil (Fisher Scientific), administered over five consecutive days.

### Animal Husbandry

Mice used in this study, including Rag2GFP reporter mice ^45^, were female and bred at Charles River UK Ltd and the Comparative Biology Unit, Royal Free Hospital. Ki67 reporter mice were maintained aged 8-12 weeks; busulfan chimeras followed age ranges indicated in the corresponding figure legends. All experimental procedures adhered strictly to UCL Animal Welfare and Ethical Review Body and Home Office regulations.

### Flow Cytometry Analysis

#### Sample preparation

Flow cytometric analyses were performed on a minimum of 2 × 10^6^cells isolated from the organs of interest. Cells were stained with monoclonal antibodies (mAbs) in PBS (100*µL*) at a saturating concentration and for one hour in the dark at 4^*°*^C.

#### Immunophenotyping

The panel detected various surface antigens using commercially available mAbs, including B220 (BioLegend), CD21 (c; Biolegend), CD93 (Biolegend), CD95 (BD Biosciences), IgD (Biolegend), and IgM (eBioscience). Live/dead staining utilized a Near-IR dye (Life Technologies). A secondary staining step was performed using either streptavidin-BUV395 (BD Biosciences, 0.5 *µ*g/ml) or streptavidin-PerCP-Cy5.5 (BioLegend, 0.4 *µ*g/ml); cells were incubated with the secondary stain for 30 minutes at 4°C in the dark. Following a thorough wash in handling media, samples were immediately analyzed by flow cytometry.

#### Intracellular staining

For intracellular markers, cell fixation and permeabilization were executed using the FoxP3 transcription factor staining buffer set (eBioscience). Proliferation was assessed via detection of Ki67 using SolA15-FITC or SolA15-PE (eBioscience).

#### Data acquisition

All data were processed using FlowJo v10 (Becton Dickinson and Company).

### Mechanistic and Statistical Modeling

Mathematical models, schematically illustrated in **Figure 3A** and described in detail in **SI section S2** were simultaneously fitted to the time courses of total cell numbers, normalized chimerism levels, and Ki67 expression in both host and donor populations. These fits incorporated empirical descriptions of the candidate precursor populations (detailed in **SI Section S1**).

Statistical analysis utilized a Bayesian estimation approach, described in **SI section S3**. The joint likelihood of all observations was formulated, which was then combined with prior distributions for model parameters to generate posterior probability distributions. This procedure provided the Leave-One-Out Information Criterion (LOO-IC), serving as both a metric for goodness-of-fit and an indicator of model complexity. Model fits presented in **figures 3** and 4 were generated using independent samples drawn from these estimated posteriors. Details regarding the modeling of MZ B cell dynamics in young mice are provided in SI **section S5**.

## ACKNOWLEDGEMENTS

The authors acknowledge financial support from the United Kingdom Medical Research Council (MR/P011225/1 to B.S.) and the National Institutes of Health (R01 AI170965 to S.R, R01 AI093870 to A.J.Y).

## Supplementary Information

### Supplementary Note S1: Modeling dynamics of precursor influx into MZ B pool

We assessed four candidate precursor populations — T1, T2, follicular B (Fo B), and marginal zone precursor (MZP) B cells — as potential sources of MZ B cells. Our modeling framework was constrained by empirical data, including absolute cell counts, donor chimerism within each source, and the fraction of Ki67^+^ donor and host cells in the precursor populations.

#### Total size of the precursor pool

We observed that the absolute number of cells in each precursor population remained stable across the mouse lifespan. Therefore, we defined the precursor pool size, denoted as *P*, as the mean value of the observed counts. Descriptions of each precursor population size are shown in **figure S1A**.

#### Chimerism in precursor populations

We described the changes in the fraction of donor-derived cells within these precursors using the function,

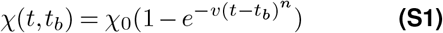

where *t* is the animal age and *t*_*b*_ is the age at bone marrow transplantation (BMT). The parameters *χ*_0_ (stable chimerism) and *v* (rate of approach to stability) were estimated by fitting **equation S1** to the observed time course of donor chimerism using non-linear least squares. We treated *n* as a fixed parameter and tested four different values (*n* = 0.5, 1, 1.5, and 2) across all subsets. The donor fraction kinetics in T1 and T2 B cells were best captured by *n* = 1, whereas those in Fo B and MZP cells were best described by *n* = 2. Model fits and parameter estimates for each precursor are shown in **figure S1B**.

#### Ki67 expression dynamics in the precursors

The fraction of Ki67 expression in both donor and host derived precursor cells was described using empirical functions. For transitional B cells, the Ki67 expression dynamics were modeled as,

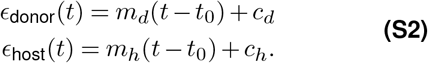

where *t* is the animal age and *t*_0_ = 40 days is the age of the youngest animal at BMT in the adult dataset. Ki67 expression kinetics for Fo B and MZP differed from transitional B cells, with donor-derived cells modeled as an exponential decay:

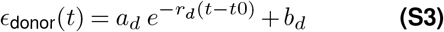

where *t* is the animal age and *t*_0_ = 40 days.

The fraction of Ki67^+^ host-derived Fo B and MZP cells remained relatively constant over time. Therefore, *ϵ*_host_ was assumed to be constant and set to the median observed values (0.068 and 0.078, respectively). Parameters (*m*_*d*_, *c*_*d*_, *m*_*h*_, *c*_*h*_, *a*_*d*_, *r*_*d*_, *b*_*d*_) were estimated by fitting the **equations S2** and **S3** to the observed fractions of Ki67^+^ donor- and host derived cells using least squares. Comprehensive model fits and parameter estimates are shown in **figure S1C**.

### Supplementary Note S2: Modeling population dynamics of MZ B cells in busulfan chimeras

We employed a mechanistic modeling approach to characterize MZ B cell dynamics in busulfan chimeric mice that describes three key observables: total MZ B cell numbers over time, donor chimerism within the MZ compartment, and the temporal dynamics of Ki67 expression in host and donor subsets. By fitting the model to all three measurements simultaneously, we aimed to quantify the relative contributions of precursor influx, self-renewal, and cellular loss to MZ B cell homeostasis.

#### S2A Deriving the loss rate under steady-state condition

The dynamics of the total MZ B cell population can be described by summing the donor and host compartments and is given by,

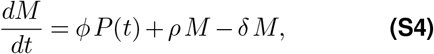

where *ϕ* denotes the per capita influx rate from the source, *ρ* is the per capita proliferation rate, and *δ* is the per capita loss rate due to death or onward differentiation. The total precursor pool size is *P* (*t*) = *P* (*t*) *χ* (*t, t*_*b*_) + *P* (*t*) (1 − *χ* (*t, t*_*b*_)).

Since MZ B cell numbers remain stable in adult mice, we assumed that the system was at steady state, such that the rate of production balances the rate of loss. Imposing this condition on **equation S4** yields:

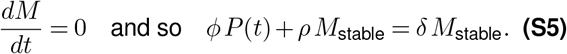

Rearranging for *δ*, we get

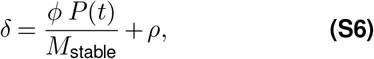

where *M*_stable_ is the steady-state total cell count. Importantly, **equation S6** reduces the number of free parameters by expressing *δ* in terms of *ϕ* and *ρ*, thereby improving parameter identifiability. The loss rate was allowed to co-vary with the influx and division rates in the time-dependent models and is given by

**Figure S1.**
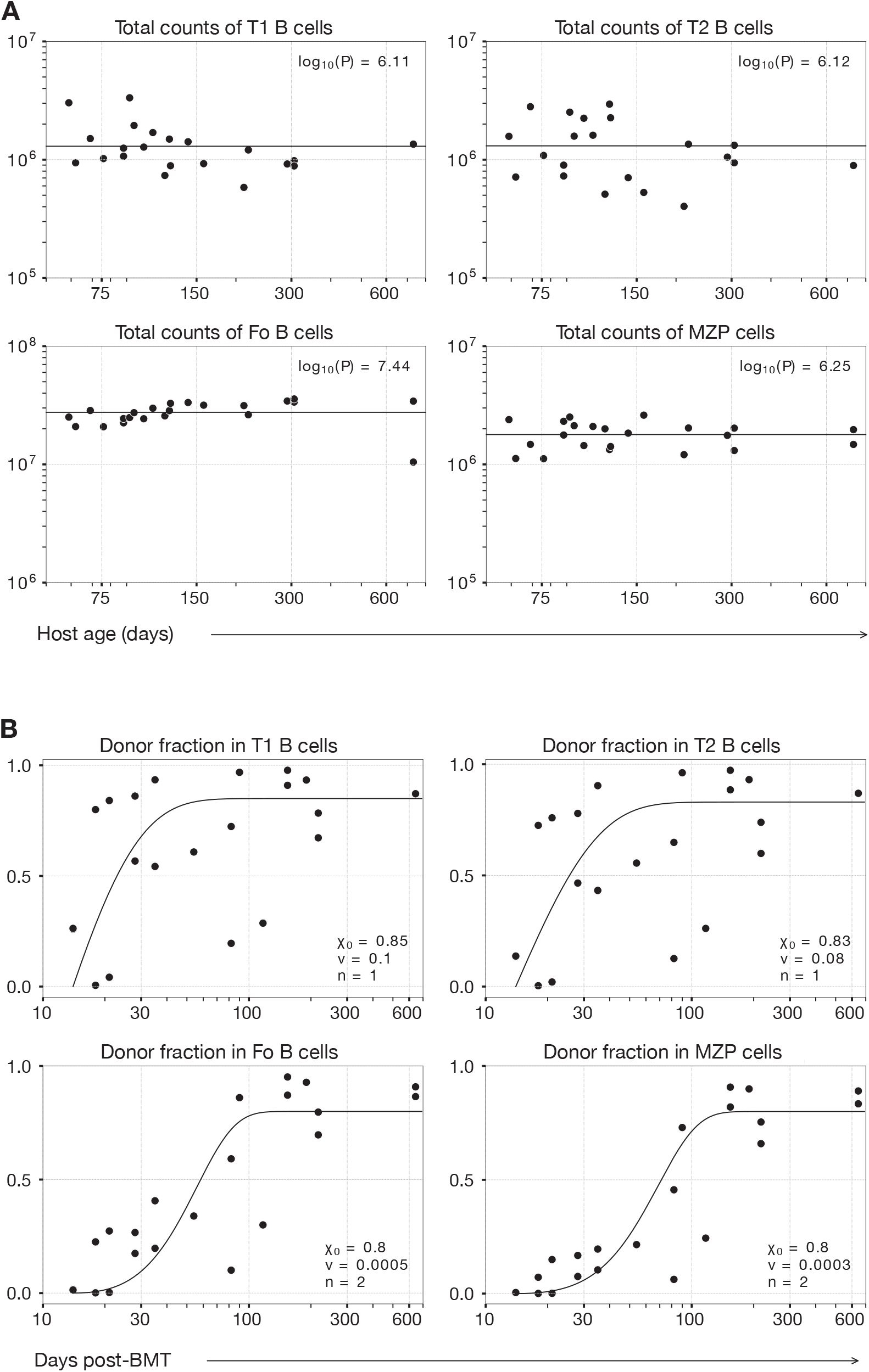
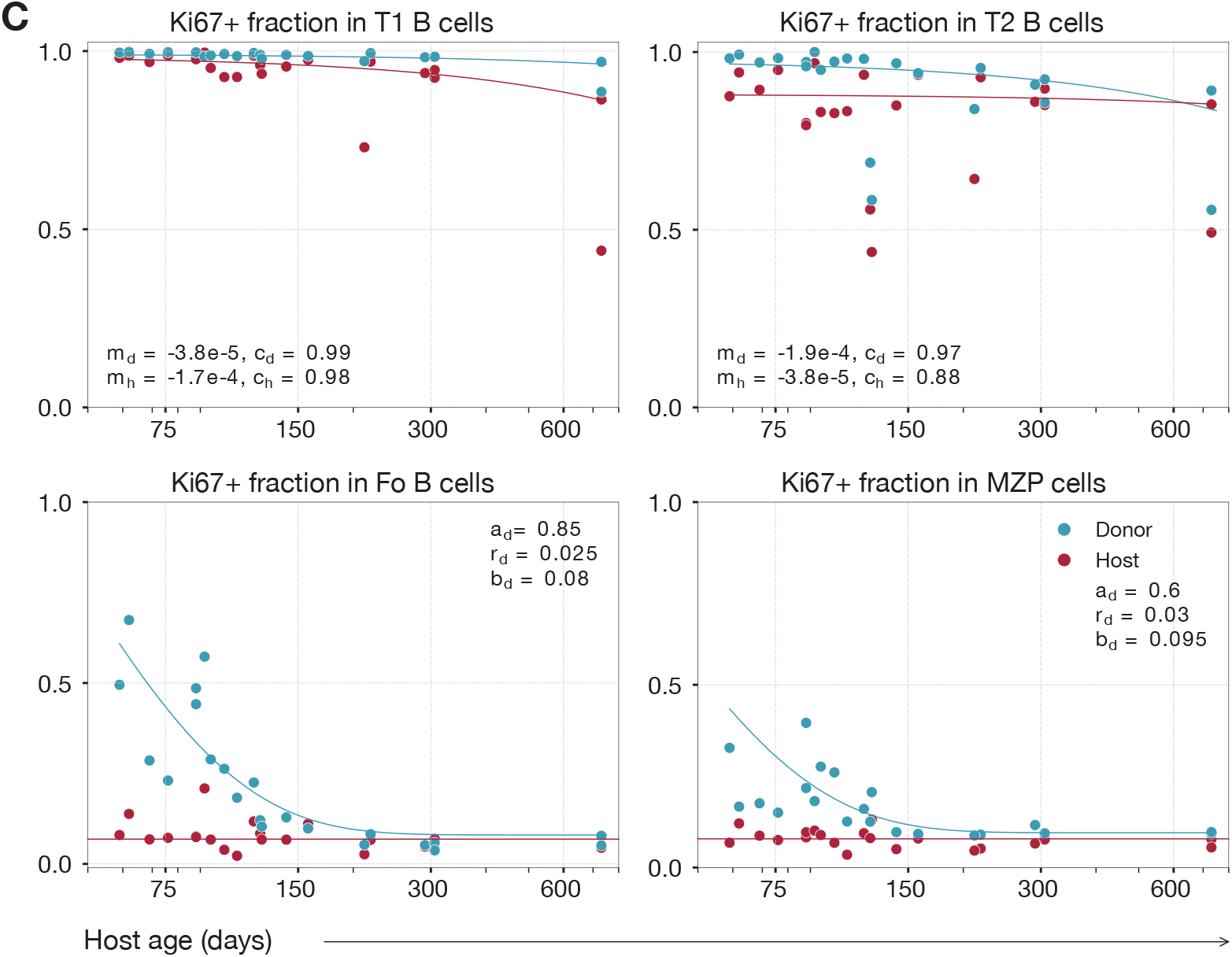
Dynamics of potential precursor populations of MZ B cells. We show the time courses of total cell counts (A), donor chimerism (B), and the proportions of Ki67^+^ donor- and host-derived precursor subsets (C). The lines in panel A show the mean estimates of the observed total counts. Similarly, the curves for donor chimerism were generated using the empirical function defined in equation S1. The Ki67^+^ fractions in donor- and host-derived T1 and T2 B-cell subsets were generated using equation S2, whereas those for donor-derived Fo B and MZP cells were generated using equation S3. For host-derived precursor Fo B and MZP cells, the Ki67^+^ fractions were fixed at the median of the observed values.

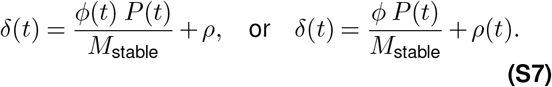

#### S2B Mathematical models for MZ B dynamics

##### Constant birth-loss model

First we considered a model that assumes that the processes of production and loss in the MZ B cell compartment remain constant over the mouse lifespan. The model was formulated as a system of ordinary differential equations (ODEs) to separately track the dynamics of Ki67^+^ and Ki67^*−*^ cells within the host and donor compartments (**equation S8**). We consider 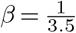 is the per capita rate of loss of Ki67 expression following mitosis.

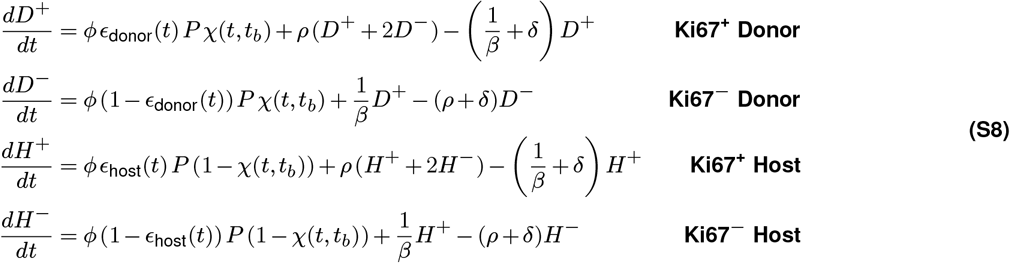

We next extended the framework to incorporate time-dependent models in order to assess the influence of host age on MZ B cell dynamics. In these models, we allowed one process at a time to vary with host age: either the per capita influx rate from the precursor population *ϕ*, or the per capita division rate *ρ*. This allows the model to capture potential age-related changes in recruitment or proliferative activity within the MZ B compartment. We tested multiple functional forms, including exponential, Hill, and sigmoid families, to explain time-varying rates.

##### Time-dependent influx model

We assumed that the per capita influx rate from the precursor population varies with host age, as described in **equation S9**. Specifically, this time dependence is best described by an exponential function as *ϕ*(*t*) = *ϕe*^*−r*(*t−t*0)^, where *t*0 = 40 days is the age of the youngest animal at BMT, and *r* is a free parameter. At any given time, all cells in the population were assumed to divide at the same rate.

##### Time-dependent division model

Here we consider that the per capita division rate *ρ* varies with host age (**equation S10**). Among the functional forms, a Hill function provided the best description of this dependence, 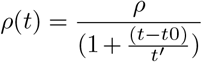 and *t*_0_ = 40 days. The parameter *t*^*′*^ represents a timescale that was inferred from the data and determines how quickly the cell division rate declines as the host ages. At any given time, all cells in the population were assumed to share the same influx rate.

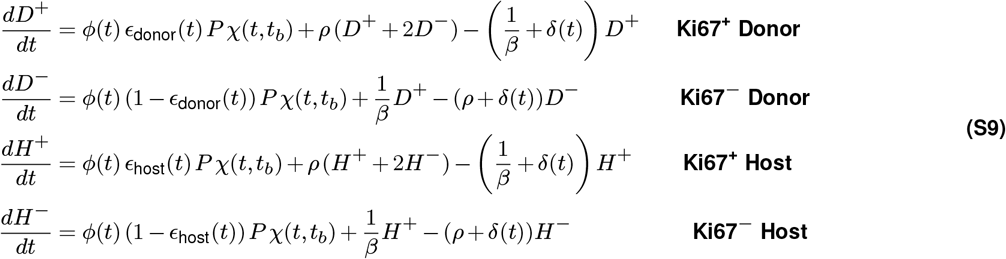

S

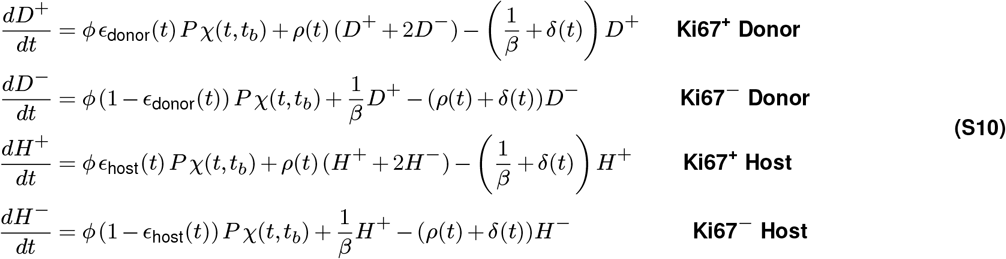

We solved these systems numerically to obtain the time courses of total cell numbers *M* (*t*), the donor fraction normalized to chimerism *f*_*d*_, and the Ki67^+^ fraction in host *κ*_*h*_ and donor *κ*_*d*_ subsets.

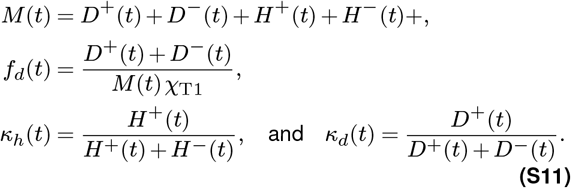

where *χ*_*T* 1_ is donor chimerism in T1, estimated from fitting **equation S1** to donor fractions in T1 B cells (**figure S1B**).

The total donor- and host-derived precursor pool sizes (*P*_donor_, *P*_host_) and their corresponding Ki67^+^ fractions *ϵ*_donor_, *ϵ*_host_ are described empirically for each precursor in **section S2**. The Ki67 loss rate *β* is fixed to 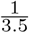 days^*−*1^ based on mean posterior estimates from our previous studies ^2,5^ modeling Ki67 turnover in T and B cell lineages. The parameters *ϕ, ρ, κ*_0_ (the Ki67^+^ fraction among host cells at the earliest age at BMT), and the steady-state MZ B cell number *M*_stable_ were estimated by fitting the model to the adult data. The time-dependent models add an additional parameter *viz. r* and *t*^*′*^ (time varying influx and division models, respectively), which were also estimated from the model fits. A comparison of the statistical support for each model is provided in **table S1**, while parameter estimates for the constant birth-loss model using transitional subsets as precursors are presented in **table S2**.

### Supplementary Note S3: Model fitting and selection criterion

Each candidate model was fitted simultaneously to four key observations measured in each busulfan chimeras *n* (*n* = 1,…, 22): total MZ B cell counts (*y*_*n*,1_), normalized donor chimerism (*y*_*n*,2_), and the fractions of Ki67^+^ cells within donor (*y*_*n*,3_) and host (*y*_*n*,4_) subsets. Parameter estimation was performed within a Bayesian statistical framework. We assumed that each observation *y*_*n,i*_ follows an independent normal distribution around its model prediction *µ*_*n,i*_:

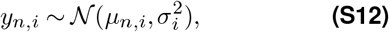

where 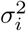 is the residual variance for the observable *i* and *µ*^*n,i*^ is the model prediction for *i*^*th*^ observation in *n*^*th*^ animal.

**Table S1.** Support for precursor:model combination from busulfan chimera data. The *ELPD* estimate measures the goodness-of-fit. We show the difference in s*ELPD* estimates (Δ *ELPD*) along with the standard error *(s*.*e*.) in its measurement. By convention, models whose Δ*ELPD* values fall within twice the *s*.*e*. are considered to have equivocal support. We find that time-varying process models fail to improve on the simplest constant birth-loss model, and models considering either T1 or T2 overall outperform models with Fo B and MZP as precursors.

| Model and $\Delta$ *ELPD* | | | |
| --- | --- | --- | --- |
| Precursor | Constant birth-loss | Time-dependent division | Time-dependent influx |
| T1 | $0 \pm 0$ | $0 \pm 0$ | $2 \pm 2$ |
| T2 | $0 \pm 0$ | $0 \pm 0$ | $2 \pm 2$ |
| FO | $12 \pm 4$ | $12 \pm 4$ | $13 \pm 5$ |
| MZP | $16 \pm 5$ | $16 \pm 5$ | $17 \pm 5$ |

**Table S2.** Parameter estimates for the constant birth-loss model. Posterior mean and 95% credible interval are shown for parameters estimated from the best-fitted constant birth-loss model considering either T1 or T2 as a precursor. The 95% credible intervals are calculated from the 2.5th and 97.5th percentiles of the posterior distributions.

| Parameter estimates from constant birth-loss model<br>(mean and 95% credible interval) |  |  |  |
| --- | --- | --- | --- |
| Parameter | Description | T1 precursor | T2 precursor |
| $\phi$ | Per-capita influx rate ($\times 10^{-2}$) | 1.9 (1.4, 2.4) | 1.9 (1.4, 2.5) |
| $\rho$ | Per-capita division rate ($\times 10^{-2}$) | 2.5 (1.9, 3.2) | 2.5 (1.9, 3.2) |
| $\delta$ | Per-capita loss rate ($\times 10^{-2}$) | 3.5 (2.9, 4.2) | 3.5 (2.9, 4.2) |

The model predictions, *µ*_*n,i*_ = *f*_*i*_(*t*_*n*_, *θ*), were obtained by solving the ODE system with parameter vector ***θ***. The following observables were evaluated at time *t*_*n*_:

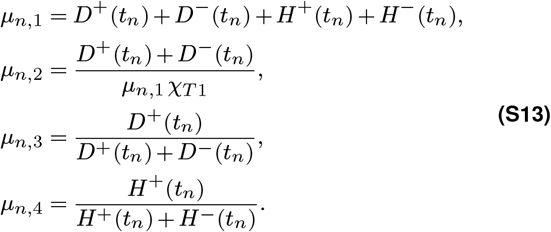

The joint log-likelihood was calculated by assuming that the four observables are conditionally independent given the parameter vector ***θ***:

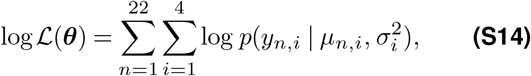

where *p*(·|.,.) is the normal probability density function defined as:

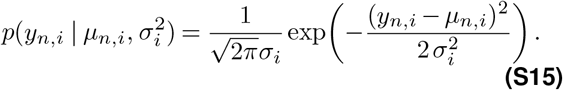

We specified weakly informative priors for all model parameters. The posterior distribution was obtained by combining the likelihood with the prior distributions using Bayes’ theorem. Posterior sampling was performed in Stan using the No-U-Turn Sampler (NUTS), an adaptive form of the Hamiltonian Monte Carlo (HMC) algorithm ^1^.

#### Model selection criterion

The expected log point-wise predictive density (*ELPD*) was adopted as the model selection criterion for the purpose of quantifying their relative statistical support. The *ELPD* was estimated for each model *M*_*j*_ by means of leave-one-out (LOO) cross-validation. The leave-one-out predictive densities were approximated from a single full-data fit by Pareto-smoothed importance sampling (PSIS-LOO; arviz.loo/compare) ^3,4^, giving

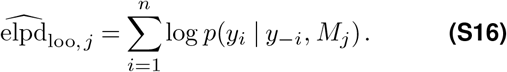

For the purpose of model comparison, the difference in *ELPD* (Δ*ELPD*) relative to the best-performing model was reported alongside the standard error of this difference. By convention, two models were considered to have statistically indistinguishable support when their Δ*ELPD* values were found to fall within twice the standard error.

**Figure S2.**
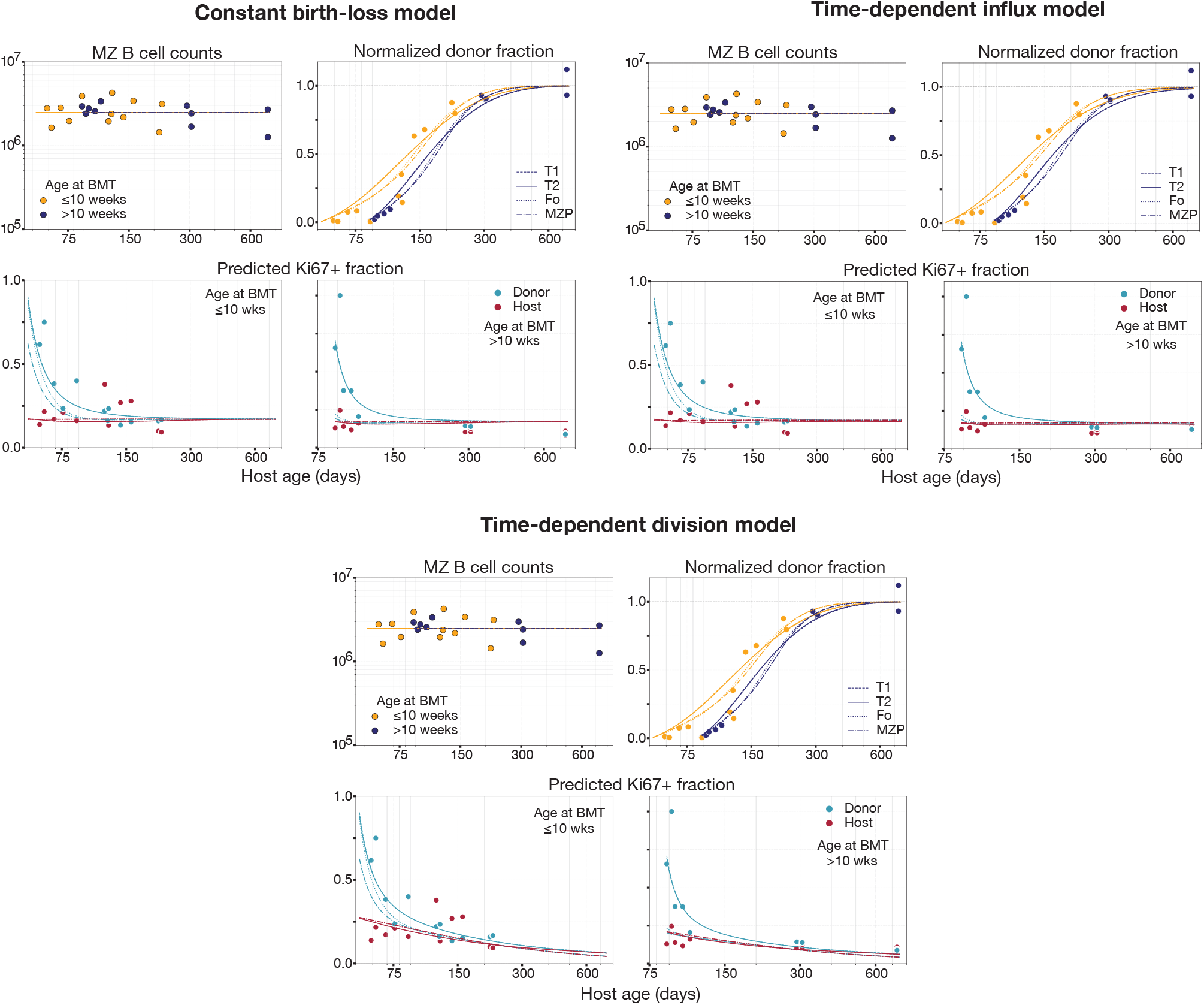
Fitted alternative models to the population dynamics of MZ B cells in busulfan chimeras. Constant birth–loss, time-dependent influx, and time-dependent division models were fitted to total MZ B cell counts, normalized donor chimerism, and Ki67 expression stratified by host and donor cells. The four fitted lines correspond to the four precursor populations considered in the analysis. For total cell counts and normalized chimerism, line colors indicate mice grouped by age at BMT: *≤* 10 weeks and *>* 10 weeks.

### Supplementary Note S4: Estimating time to near-complete replacement

We model the donor-derived fraction in the MZ compartment with

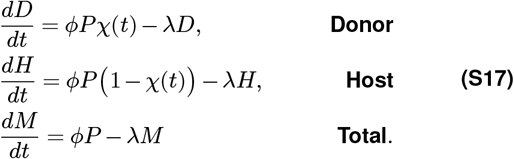

Here, *M* = *D* + *H* is the total MZ population, *P* is the source population, *ϕ* is the influx rate into *M*, and *λ* is the loss rate from *M*. The donor fraction in the source T2 subset changes over time as,

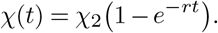

Because *M* is assumed to be at steady state, *ϕS* = *λM*. Defining the normalized donor fraction in *M* as 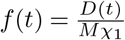, where *χ*_1_ is the donor fraction in the T1 B cells. For this particular calculation, we begin our model at the time when chimerism in the T1 subset has stabilized *i*.*e. t* = *t*_*χ*_, such that *χ*_1_ is constant.

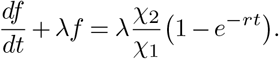

With initial condition *f* (*t*_*χ*_) = *f*_0_, the solution is,

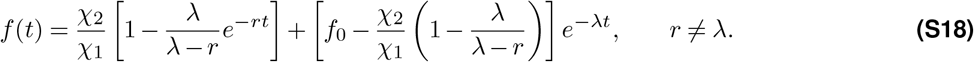

As *t* → ∞, the asymptote is

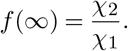

We define the replacement time, *t*_*s*_, as the first time at which *f* (*t*) reaches 95% of its asymptotic value:

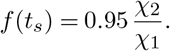

Using *f*_0_ = 0.03, and *χ*_2_*/ χ*_1_ = 0.98, so the 95% threshold is *f* (*t*_*s*_) = 0.931.

Substituting the parameter values into the solution gives

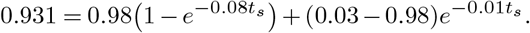

Equivalently,

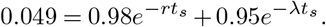

Since *λ* ≪ *r*, the slow *e*^*−λt*^ term dominates the approach to the asymptote, giving the approximation

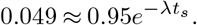

Solving for *t*_*s*_ using *λ* estimates obtained from our bestfit model yields the mean estimate of time to near-complete (95%) replacement *t*_*s*_ ≈ 300 days with lower and upper bounds of 250 and 370 days.

### Supplementary Note S5: Modeling the development of MZ B cells in young mice

#### A. Empirical descriptions of transitional B cell numbers in young mice

We treated T1 and T2 as upstream precursors, such that influx is proportional to the size of the corresponding precursor pool. The empirical descriptions of T1 and T2 precursor numbers (**figure 4A**) are

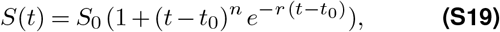

*t*_0_=10 days is the time offset used in the fit and does not represent the actual age of the animal. We fitted the equation S19 to the log-transformed cell counts and estimated *S*_0_, *n* and *r* using the least squares method.

#### B. Models with age-dependent dynamics

We found that the best-fitting constant birth–loss model fitted to the chimera data failed to capture the early developmental dynamics of MZ B cells, as it overestimated cell numbers during early life (**figure 4B**). This discrepancy suggests that the processes regulating MZ B cell dynamics differ between early life and adulthood.

To test whether developmental dynamics could be explained by age-dependent changes in a single kinetic process, we extended the model to allow influx, division, or loss to vary with host age. We considered a general age-dependent model:

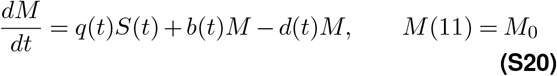

where *M*_0_=12058 cells is the observed MZ B cell count at day 11, and q(t), b(t), and d(t) denote the potentially age-varying influx, division, and loss rates respectively. In each sub-model, only one of these three processes was allowed to vary with host age while the other two were kept constant.

We considered multiple candidate functional forms, including exponential, Hill, and sigmoid families, to explain each process. The best function was selected based on simultaneous fits to total MZ B cell numbers (days 11–731), normalized donor chimerism, and proportion of Ki67^+^ within MZ B compartment in adult busulfan chimeras (from day 40 onwards), considering only the transitional B cell subsets as candidate precursors. Posterior distributions of the adult parameters were used as informative priors when fitting the developmental dynamics of MZ B cells. For the influx rate, a sigmoid function of the form:

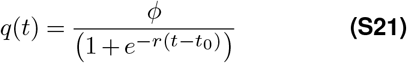

provided the best description, capturing the gradual rise in precursor recruitment from near-zero at birth toward the adult influx rate *ϕ*, with parameters *r* and *t*_0_ determining the rate and timing of this transition. The division rate was best governed by a Hill function:

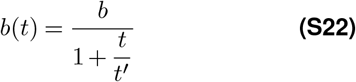

where *b* is the maximum division rate at the youngest observed age and *t*^*′*^ is the timescale over which the rate declines toward its adult value. The loss rate was best represented by an exponential decay:

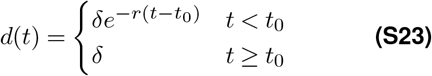

with *r* and *t*_0_ governing the rate and timing of convergence to the adult loss rate *δ*. We found that none of the models could adequately capture both the increase in cell numbers in neonates and the maintenance of stable counts in adulthood. The fits from each model are provided in **figure S3**.

#### C. Models with density-dependent processes

We observed that the population dynamics of MZ B cells follow a similar pattern to a logistic growth curve. Therefore, we considered models in which the either rate of influx or division or loss rate was allowed to vary as functions of the size of the total MZ B cell population. The general formulation for this class of model is:

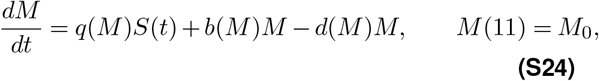

where the processes other than the varying with density are assumed to be constant.

We defined density-dependence in influx and division rates using,

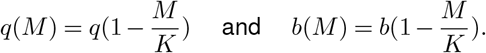

For the loss rate we considered an increasing function: 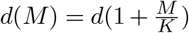. Parameter interpretations and constrains are defined below.

In all three models, the parameters *q, b, d*, and *K* were estimated by fitting the models simultaneously to the full dataset. The density-dependent influx model failed to capture the overall dynamics of MZ B cells (**figure S4**). However, while the density-dependent division with a constant influx rate described MZ B cell numbers and Ki67 expression in donor and host subsets reasonably well, they failed to adequately reproduce the normalized donor fraction dynamics in adult animals (**figure S4**).

We therefore extended the density-dependent division and loss models by allowing the influx rate to vary with host age and fitted these models to the full dataset. We explored several forms of time-varying influx rate, as described earlier. Influx varying with time sigmoidally given by the hill function as, 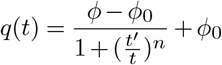reconciled the dynamics of MZ B cells across early development and homeostasis later. This model produced smooth trajectories of total pool size, *f*_*d*_ and Ki67 fractions among donor and host compartments that agreed well with the observed data. Here, *ϕ*_0_ represents the influx rate at 11 days of age, *ϕ* is the stabilized level of influx rate in adult mice, and *t*^*′*^ is the timescale at which the influx rate reaches half of its adult value. We explored different values for the hill exponent and found that *n* = 7 gives the best-fit. Rest all parameters were estimated through model fitting procedure.

### Supplementary Note S6: Quantifying MZ B cell production and clonal half-life

Using the density-dependent division model with time-varying influx, we quantified the daily production of MZ B cells as the combined contribution of precursor influx and self-renewal through proliferation (**figure 5D**). Total daily production was defined as *q*(*t*)*P* (*t*) + *b*(*M*)*M*.

We next estimated the relative contributions of precursor influx and self-renewal to total MZ B cell production (**figure 5E**). The contribution of precursor influx was calculated as

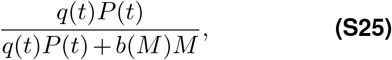

whereas the contribution of self-renewal was calculated as

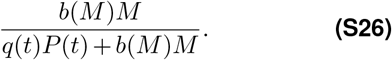

These quantities represent the fractions of total daily MZ B cell production arising from precursor recruitment and self-renewal, respectively.

Fitting the density-dependent division model with time-varying influx across the full lifespan revealed two distinct kinetic regimes, determined by the balance between cell division (*b*) and loss (*d*). In young animals, division exceeds loss (*b > d*), resulting in a positive net growth rate and expansion of the MZ B cell compartment. This accumulation of MZ B cells during the neonatal period is characterized by a population doubling time of 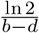. In adult animals (4 months and older) the balance reverses (*d > b*), and the same framework predicts the clonal half-life, 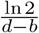. Similarly, we estimate the clonal lifetime as the inverse of the net loss rate 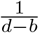.

### Supplementary Note S7: Carrying capacity in a density-dependent growth model with influx and loss

In modeling the population dynamics of MZ B cells, we analyze a scalar ordinary differential equation that extends the classical logistic growth model by incorporating a recruitment term *s* and a linear loss rate *d*. In the classical logistic framework ^6^, the crowding parameter *K* serves a dual role. It governs density-dependence and simultaneously defines the equilibrium population size, *i*.*e*. the carrying capacity. This equivalence breaks down, however, once immigration and efflux are introduced. Here, we show that the carrying capacity, *i*.*e*. the stable population size, is not defined by *K* but rather emerges from the interplay between immigration, division, and loss.

Let *M* (*t*) ≥ 0 denote the size of a population at time *t* ≥ 0. We consider the following initial-value problem:

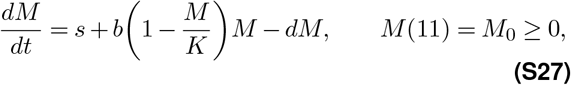

where the parameters are non-negative real numbers with the following interpretations:

The term *b*(1 − *M/K*) is the classical logistic proliferation rate: division occurs at the maximal rate *b* when the population is sparse and is completely suppressed when *M* = *K*. **Equation S27** augments this with a source term *s* and a first-order loss term *dM*.

#### Steady-state analysis

##### Equilibrium Equation

Setting *dM/dt* = 0 in Eq. (S27) and collecting terms yields the quadratic:

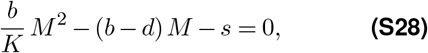

whose roots are

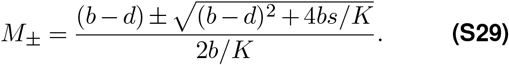

The discriminant Δ = (*b* − *d*)^2^ + 4*bs/K* is strictly positive for all admissible parameters, so both roots are real and distinct. Since the product of the two roots equals − *sK/b <* 0, one root is necessarily positive and the other negative. Only *M*_+_ is biologically meaningful.

Therefore, the carrying capacity of system Eq. (S27) is the unique positive equilibrium *M*^*∗*^, defined as

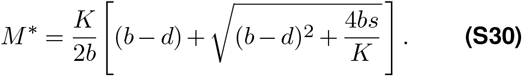

It is the stable population size to which every trajectory with *M*_0_ ≥ 0 converges.

The parameter *K* is *not* the carrying capacity as described by **equation S27**. Rather, *K* is the value of *M* at which the density-dependent proliferation term *b*(1 − *M/K*)*M* vanishes. Only in the degenerate case *s* = *d* = 0 does the system reduce to the classical logistic equation, for which *M*^*∗*^ = *K*.

##### Comparison with the Classical Logistic Model

To make explicit how *s* and *d* shift the equilibrium away from *K*, we compare them across cases, *viz*., Classic, Loss only, influx only, and general scenario where both influx and loss are involved.

| Conditions | Equilibrium | Relation to $K$ |
| --- | --- | --- |
| $s = 0, d = 0$ | $M^* = K$ | $M^* = K$ |
| $s = 0, d > 0$ | $M^* = K(1 - d/b)$ | $M^* < K$ |
| $s > 0, d = 0$ | $M^* > K$ (if $s$ large) | $M^* > K$ |
| $s > 0, d > 0$ | Eq. (S30) | depends on $s$ and $d$ |

In the classical logistic framework, the carrying capacity *K* plays a dual role: it is simultaneously the saturation parameter for density-dependent growth *and* the long-run equilibrium population size. This coincidence disappears as soon as additional biological processes — loss or immigration — are introduced.

In this model, *K* retains its mechanistic interpretation: it is the population size at which cellular competition completely suppresses new divisions. However, the system does not equilibrate at *K* because the influx term continues to add individuals even when *M ≥ K*, and the loss term continues to remove them for all *M >* 0. The true carrying capacity *M*^*∗*^ emerges from the balance of all three forces — proliferation, influx, and loss — and is given by **equation S30**.

Model comparison across all models is shown in (**table S3**). We observed comparable support density-dependent division and loss models with varying influx rates. We show the parameter estimates for the best-supported model in **table S4**.

---

**Figure S3.**
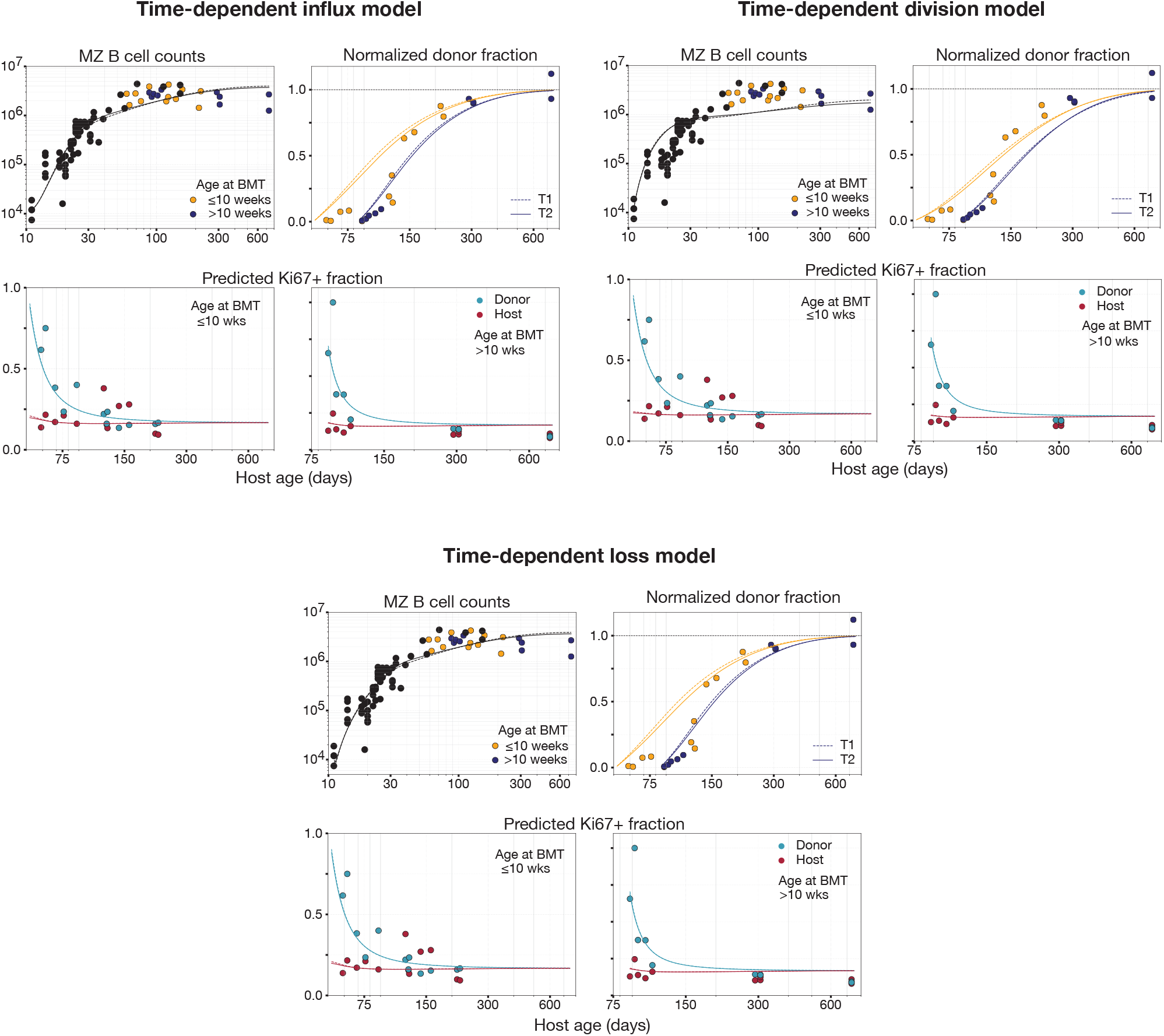
Fitted alternative time-dependent models to the developmental dynamics of MZ B cells. We tested models in which either the influx rate from the precursor population, the rate of self-renewal, or the loss rate of MZ B cells was allowed to vary with mouse age. Models were fitted to MZ B cell counts in young mice and busulfan chimeras, as well as to normalized donor chimerism and Ki67^+^ frequencies among donor and host cells in chimeric mice, assuming transitional B cells as the precursor population. None of these models adequately captured the rapid increase and subsequent establishment of stable MZ B cell numbers.

**Table S3.** Comparison of models describing MZ B cell dynamics using data from both young and busulfan chimeric mice. We showed the difference in *ELPD* estimates (Δ*ELPD*), the absolute difference between model’s *ELPD* estimate and the highest *ELPD*, along with its standard error (*s*.*e*.. Density-dependent models with variable influx (lowest Δ*ELPD*) received the strongest support from the data.

| Source | Model and $\Delta ELPD$ | | | | | | | |
| --- | --- | --- | --- | --- | --- | --- | --- | --- |
|  | Time-dependent |  |  | Density-dependent |  |  | Density-dependent (vary influx) |  |
|  | Influx | Division | Loss | Influx | Division | Loss | Division | Loss |
| T1 | 16 ± 4 | 51 ± 8 | 15 ± 4 | 49 ± 8 | 30 ± 8 | 41 ± 8 | 0 ± 1 | 2 ± 1 |
| T2 | 14 ± 3 | 52 ± 8 | 12 ± 3 | 50 ± 8 | 30 ± 8 | 41 ± 8 | 0 ± 0 | 3 ± 2 |

**Figure S4.**
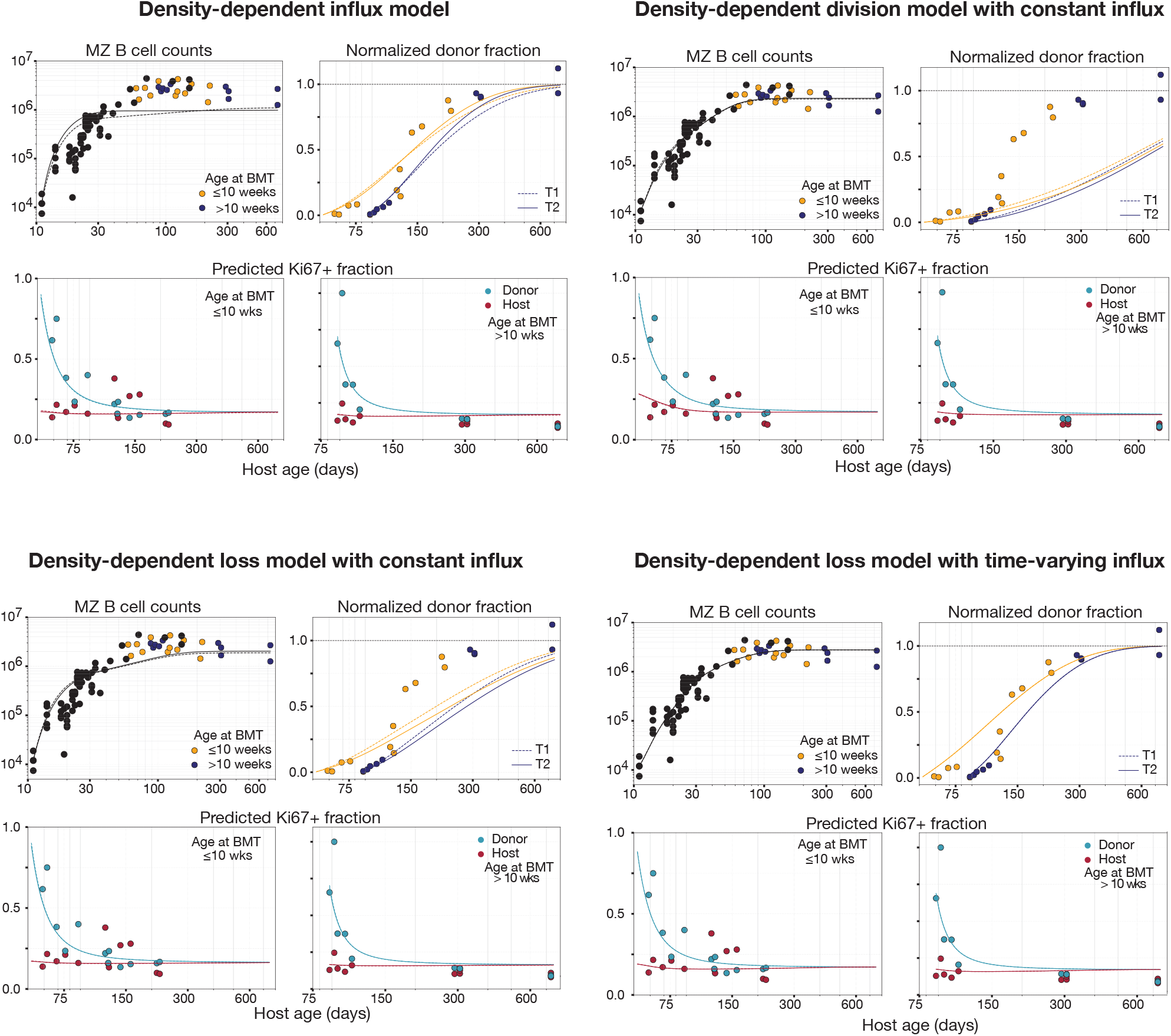
Fitted alternative density-dependent models to the developmental dynamics of MZ B cells. We fitted density-dependent models to MZ B cell numbers in young and chimeric mice, as well as to normalized donor fraction and Ki67^+^ frequencies among donor- and host-derived MZ B cells in chimeras, assuming transitional B cells as precursors.

**Table S4.**
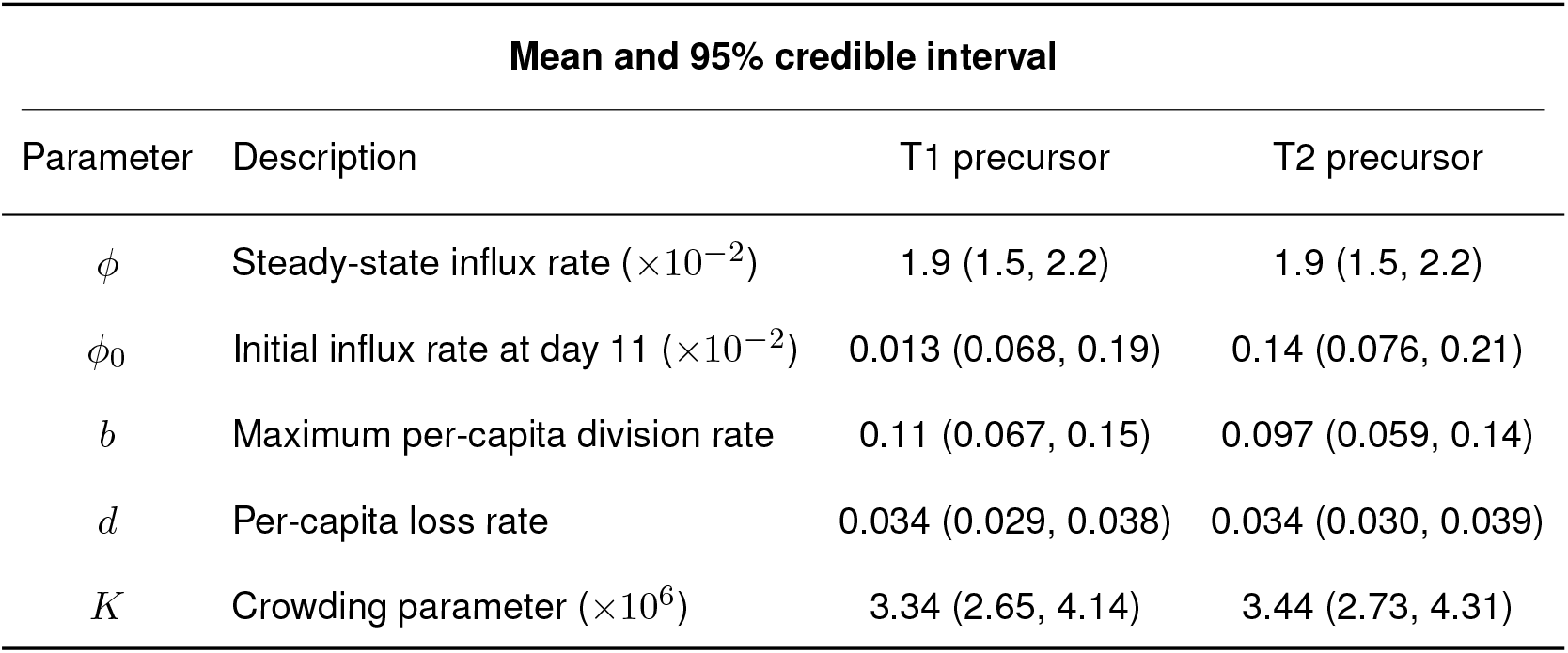
Parameter estimates for the best-supported density-dependent division model with time-varying influx.

